# When two’s a crowd - Structural mapping of PEAK pseudokinase interactions identifies 14-3-3 as a molecular switch for PEAK3/Crk signaling

**DOI:** 10.1101/2022.09.01.506260

**Authors:** Michael J. Roy, Minglyanna G. Surudoi, Ashleigh Kropp, Jianmei Hou, Weiwen Dai, Joshua M. Hardy, Lung-Yu Liang, Thomas R. Cotton, Bernhard C. Lechtenberg, Toby A. Dite, Xiuquan Ma, Roger J. Daly, Onisha Patel, Isabelle S. Lucet

## Abstract

PEAK pseudokinases regulate cell migration, invasion and proliferation by recruiting key signaling proteins to the cytoskeleton. Despite lacking catalytic activity, alteration in their expression level is associated with several aggressive cancers. Here, we elucidate new molecular details of key PEAK signaling interactions with the adapter proteins CrkII and Grb2 and the scaffold protein 14-3-3. Our findings rationalize why the dimerization of PEAK proteins has a crucial function in signal transduction and provide biophysical and structural data to unravel binding specificity within the PEAK interactome. We identify a conserved high affinity 14-3-3 motif on PEAK3 and demonstrate its role as a molecular switch to regulate CrkII binding. Together, our studies provide a detailed structural snapshot of PEAK interaction networks and further elucidate how PEAK proteins, especially PEAK3, act as dynamic scaffolds that exploit adapter proteins to control signal transduction in cell growth/motility and cancer.

## INTRODUCTION

The pseudopodium enriched atypical kinase (PEAK) family of proteins, which comprises Sugen kinase 269 (Pseudopodium-enriched atypical kinase 1, PEAK1)^1^, SgK223 (PEAK2), an ortholog of rat Pragmin^2^ and mouse Notch activation complex kinase (NACK)^3^, and the recently identified PEAK3 (C19orf35)^4^, play critical roles in actin cytoskeleton remodeling, influencing cell migration and invasion in normal and cancer cells^5-8^. Abnormal expression of PEAK proteins alters cell morphology and confers enhanced migratory and invasive characteristics to cells^1,9-12^, implying a role of PEAKs in the spatio- and temporal assembly of signaling hubs at focal adhesions (FAs) and with the actin cytoskeleton ^4,13-16^.

PEAK proteins are a unique group of pseudokinase (PsK) scaffolds that can self-assemble regulatory networks. Their scaffolding activities rely on dimerization, post-translational modifications and recruitment of modular adapter proteins. PEAK family proteins homo- and/or heterodimerize via a unique alpha-helical domain, the split helical dimerization (SHED) domain which flanks the PsK domain (Fig. 1), as recently elucidated by the structures of PEAK2^17,18^ and PEAK1^19^.

**Fig. 1:**
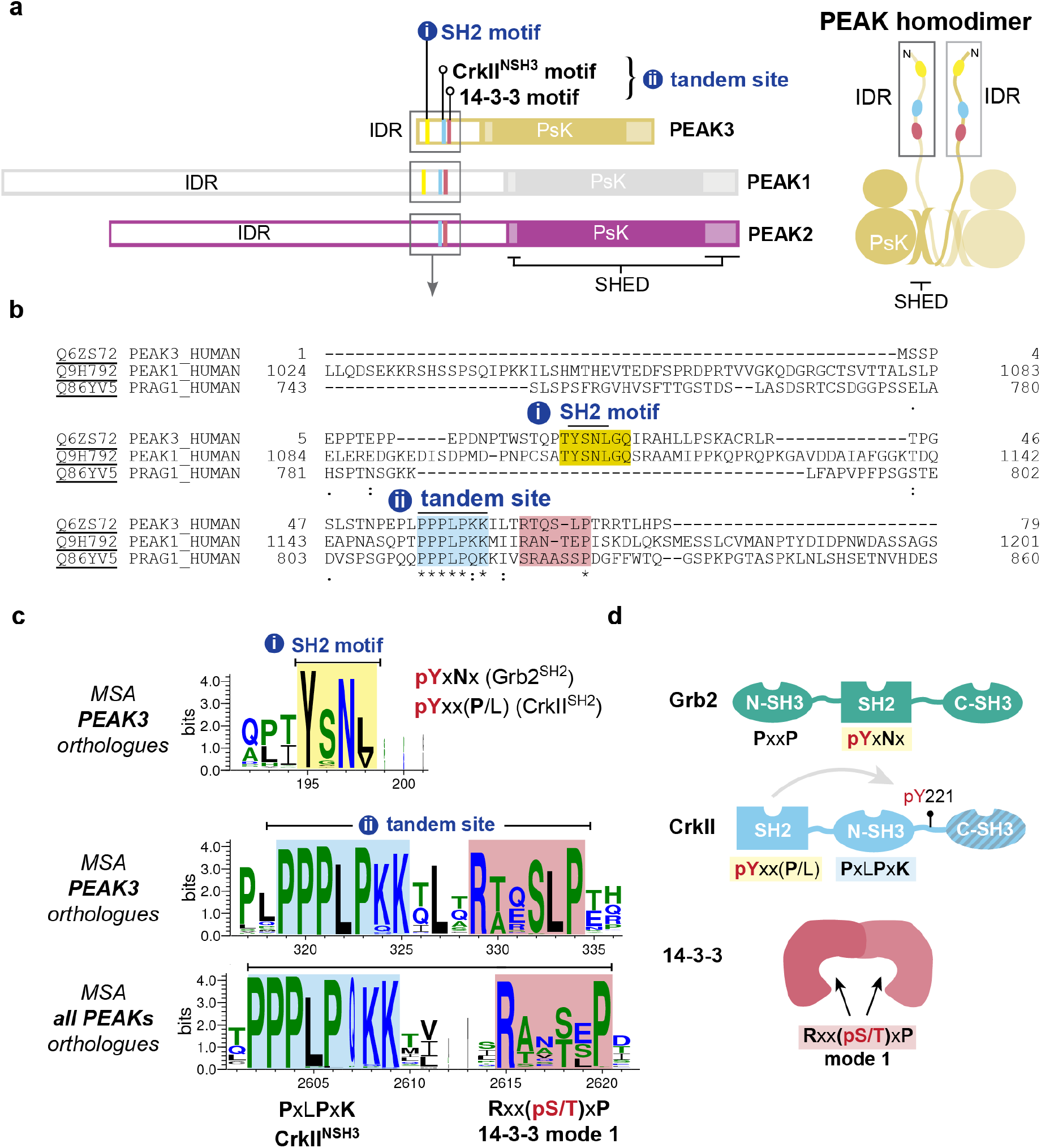
PEAK domain organization and interaction motifs in N-terminal IDR. **a**, Domain organization of the PEAK family and diagram of PEAK homodimer arrangement showing SHED, PsK and IDR motifs identified (boxed). **b**, Sequence alignment of the N-terminal IDR of human PEAK3 with the corresponding region of PEAK1 and PEAK2. **c**, Multiple Sequence Alignment (MSA) of PEAK vertebrate orthologues highlighting Short Linear Interaction Motifs (SLiMs) identified in retions of high sequence sonservation, including a pY/SH2 motif (Grb2^SH2^/CrkII^SH2^) and the tandem site encompassing a proline rich motif (CrkII^NSH3^) and conserved putative 14-3-3 motif. **d**, Schematic showing overall domain organization of Grb2, CrkII and 14-3-3, highlighting sequence motifs.

In addition to the conserved PsK and SHED domains, all PEAK proteins have an N-terminal intrinsically disordered region (IDR) (Fig. 1) replete with short linear motifs (SLiMs) – conserved sequence motifs that can that form dynamically regulated docking sites to recruit interactors involved in PEAK signalling. To date, these IDRs have only been partially characterized^4,14,17-20^. Several tyrosine phosphorylation sites have been identified within the large IDRs of PEAK1 and PEAK2 (IDR length ∼1200 and ∼900 residues, respectively) that regulate cell proliferation, migration and invasion, and in some cases the recruited proteins with Src Homology 2 (SH2) and phosphotyrosine binding (PTB) domains have been characterized. In PEAK1, Tyr^665^ is phosphorylated by Src, linking PEAK1 to the Src-p130Cas-Crk-paxillin pathway that regulates FA dynamics^21^. Additionally, phosphorylation of Tyr^635^ by Lyn creates a binding site for the SH2 domain of the adapter protein Grb2, which leads to the activation of the Ras/Raf/Erk signaling pathway that controls cell proliferation and invasion^9^. Both PEAK1 and PEAK2 are linked to the signaling adapter Shc1^7,22^ and share a conserved ‘EPIYA’ phosphotyrosine motif (PEAK2 Y^411^; PEAK1 Y^616^), which serves as a docking site for the SH2 domain of the C-terminal Src kinase (Csk). This interaction brings Csk to FAs where it likely regulates cell morphology and cell motility by modulating the activity of Src family kinases (SFKs)^16^.

In contrast, the role of PEAK3 in signaling is poorly understood. PEAK3 has a more restricted expression profile than PEAK1/PEAK2; predominantly expressed in granulocytes and monocytes (Human Protein Atlas, proteinatlas.org)^23^ suggesting a specialized function^8^. PEAK3’s IDR is significantly smaller than PEAK1/PEAK2 (PEAK3 IDR length ∼130 residues), but still harbors various SLiMs, some of which are conserved in PEAK1 and PEAK2. Almost all reported cellular interactions of PEAK3 with binding partners appear to require PEAK3 dimerization^4,14,23^. Amongst PEAK3 interactors identified from proteomic/cellular studies are the adapter proteins CrkII/CrkL^24^. CrkII/CrkL are highly similar but non-identical proteins, with roles in focal adhesion signaling, cell migration, and cancer^24^. They both share the identical domain architecture (SH2-NSH3-CSH3) (Fig. 1d) and identical binding preferences in SH2 and NSH3 domains but are reported to have a distinct structural architecture and interactome^24^. PEAK proteins have been shown to interact with CrkII via a conserved proline-rich motif (PRM) present in the N-terminal IDR of all PEAK proteins that binds the CrkII NSH3 domain (CrkII^NSH3^; consensus sequence PxLPxK) (Fig. 1d)^4,23^. Additionally, for PEAK3, and likely also PEAK1/2, CrkII binding requires PsK dimerization via the SHED domain and is impacted by mutations which disrupt the PsK domain conformation and dimerization^4,14^. Beyond this, two recent studies have identified further PEAK3 interactors, including: adapter protein Grb2; the E3 ubiquitin ligase Cbl; proline-rich tyrosine kinase 2 (Pyk2); and Arf GTPase-activating protein 1 (ASAP1)^14,23^. Grb2 (domain organization NSH3-SH2-CSH3) is involved in multiple cellular signal transduction pathways, most prominently downstream of the epidermal growth factor (EGF) receptor to the mitogen-activated protein kinase (MAPK) signalling cascade (via Grb2/Sos complex)^25^ but also survival (phosphatidylinositol 3-kinase (PI3K)/AKT) signaling (via Grb2-associated binder 1, Gab1)^26^. Other PEAK3 interactors identified include 14-3-3 proteins^4^, ubiquitous dimeric regulatory/scaffold proteins, of which there are seven isoforms in humans (β, γ, ε, η, σ, τ, and ζ). 14-3-3 proteins^27,28^ recognize specific phosphoserine/threonine motifs in partner proteins, including kinases, often with regulatory roles in cellular pathways, such as to alter conformation, activity or sub-cellular localization^29-31^.

In this study, we use an integrated bioinformatic, biochemical, and structural biology approach to further characterize the interactions and scaffolding activity of PEAK proteins. We identify highly conserved protein docking motifs within the N-terminal IDR of PEAK proteins and structurally characterize the phospho-dependent interaction of PEAK1/PEAK3 with the adapter protein Grb2. We provide new molecular details of PEAK interactions with CrkII and identify a role for the scaffold protein 14-3-3 as a new regulator of PEAK3 signaling. Lastly, we demonstrate that phosphorylation at PEAK3 S69 generates a high affinity 14-3-3 binding site. Binding of 14-3-3 to PEAK3 pS69 generates a highly stable PEAK3:14-3-3 dimer:dimer resulting in markedly reduced binding of CrkII to PEAK3 dimers. These findings contextualize why dimerization of PEAKs has a crucial function in signal transduction and demonstrate how signal specificity amongst the family is achieved. We demonstrate a 14-3-3-mediated molecular switch mechanism involving PEAK3/CrkII, providing new insights into the dynamic role of PEAKs in cell migration and invasion.

## RESULTS

### Sequence analysis of the PEAK3 N-terminal IDR identifies conserved interaction motifs

The N-terminal IDRs of PEAK family proteins contain numerous predicted SliMs; several have confirmed roles in PEAK1/2 signalling^4,9,14,21,23,32,33^, but it is likely that other important interaction sites remain to be validated, in particular those critical to PEAK3 signalling for which less is known. To further study the functionally relevant interactors of PEAK3, we first conducted an extensive bioinformatic analysis of the PEAK3 N-terminal IDR, which is comparatively shorter than those of PEAK1/2. We compared PEAK3 orthologs and direct conservation of this region with PEAK1 or PEAK2 (Fig. 1). This analysis highlighted two regions of interest. The first was a tyrosine/SH2 motif (YSNL; PEAK3^Y24^/PEAK1^Y1107^) that is highly conserved in PEAK3 and PEAK1 vertebrate orthologs (and absent in PEAK2 vertebrate orthologs) that matches the known phosphotyrosine consensus motif for the Grb2 SH2 domain (Grb2^SH2^; consensus motif: pYxNx) and the CrkII SH2 domain (CrkII^SH2^; consensus motif: pYxx(P/L), proline typically preferred) (Fig. 1a-d and Extended Data Fig. 1)^14,34^. The second region we identified was a ‘tandem site’ that comprises two directly adjacent motifs that were both conserved: a PRM known to bind CrkII^NSH3^ (consensus motif: PxLPxK)^4,14^; and a putative binding site for 14-3-3 proteins (mode 1 consensus motif: Rxx(pS/T)xP) (Fig. 1a-d). This tandem site was found to be conserved not only in vertebrate orthologs of PEAK3, but also in vertebrate orthologs across the PEAK family (Fig. 1c and Extended Data Fig. 1). While cellular studies have shown that PEAK3 interaction with Grb2 and CrkII^4,14^ are functionally important, neither have been directly characterized structurally or biophysically, nor have there been reports of a functional role for 14-3-3 with PEAK proteins.

### The conserved PEAK3^Y24^/PEAK1^Y1107^ SH2 motif is phosphorylated by Src and interacts with Grb2^SH2^

We first focused on the molecular characterization of the conserved PEAK3^Y24^/PEAK1^Y1107^ SH2 motif with Grb2. We recently identified phosphorylation of the PEAK3 TYSNL (pY24) motif in cells and showed this is required for recruitment of Grb2, and possibly CrkII and ASAP1^14^, an Arf GTPase-activating protein that regulates cytoskeletal remodeling and is associated with tumor progression and invasiveness^33,35,36^. In MCF-10A cells, phosphorylation of PEAK3 at this site is blocked by treatment with either the SFK inhibitor eCF506 or the SFK/Abl inhibitor dasatanib, suggesting that this site is dependent on SFK activity^14^. We recently also described the expression and purification of full length recombinant PEAK3 (PEAK3^FL^) and N-terminally truncated forms of PEAK1 (PEAK1^IDR1^ that includes PEAK1^Y1107^ site, residues 1082-1746) and PEAK2 (PEAK2^IDR1^, residues 802-1406) from insect cells (refer to online methods)^37^ – all of which also include the tandem motif in the IDR in addition to SHED/PsK domains. Building on this, we performed an *in vitro* kinase assay with purified recombinant PEAK3^FL^ and PEAK1^IDR1^ proteins followed by tandem mass spectrometry (MS/MS) analysis of tryptic peptides. We found that Src, but not Abl, can indeed phosphorylate this conserved SH2 motif on PEAK3^FL^ (pY24) and PEAK1^IDR1^ (pY1107) corroborating our recently reported cellular data^14^ (Fig. 2a).

**Fig. 2:**
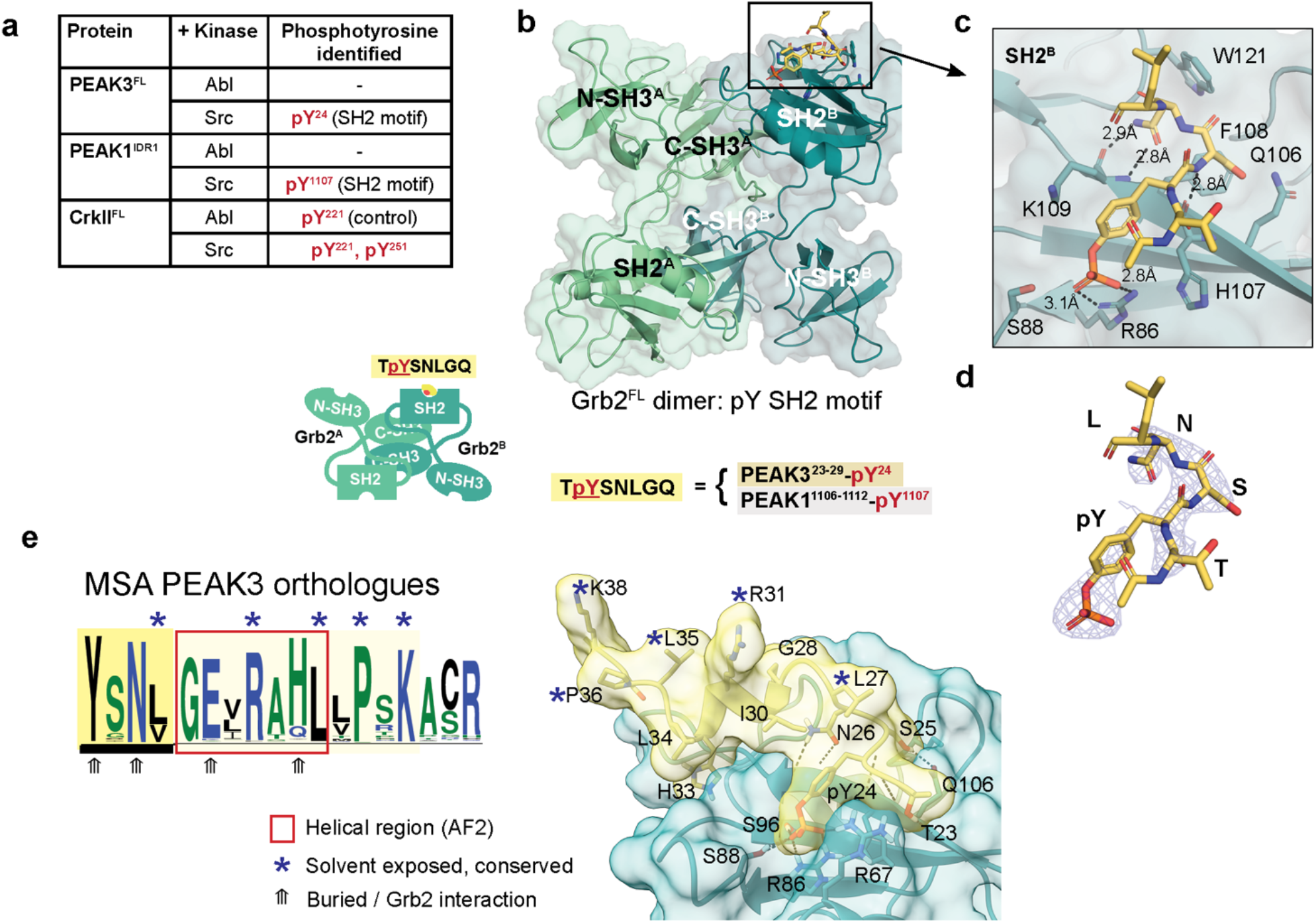
Structural and biophysical analysis of PEAK/Grb2^SH2^ interaction. **a**, TYSNL site of PEAK3^FL^ (Y24) and PEAK1^IDR1^ (Y1107) can be phosphorylated by Src (MS/MS analysis of tryptic peptides following *in vitro* kinase assay of PEAK3^FL^ and PEAK1^IDR1^ available in Source Data). Both Abl and Src can phosphorylate CrkII^Y221^, a known Abl phosphorylation site,^69^ demonstrating that both Abl and Src kinases are active. **b**, Ribbon representation of the structure of Grb2^FL^:PEAK SH2-pY peptide of PEAK3^23-29^-pY24 / PEAK1^1106-1112^-pY1107 showing dimer arrangement (chain A in light green and chain B in dark green) with cartoon illustrating domain organization. **c**, Zoom in highlighting peptide groove and the PEAK phosphopeptide (yellow) adopting a characteristic β-turn forming three important hydrogen bonds to the Grb2 backbone with residues Arg 86, His 107 and Lys109. Peptide and interacting residues of Grb2 displayed in a stick representation. **d**, Unbiased Fo-Fc omit map (grey mesh, contoured at 3.0 σ) showing peptide density prior to modelling and final modelled PEAK3/PEAK1 phosphopeptide (<u>T</u>**<u>pY</u>**<u>SNL</u>GQ, yellow sticks, underlined residues modelled). **e**, HADDOCK docked model of extended helical PEAK3^23-38^-pY24 peptide bound to Grb2, showing solvent exposed residues of high sequence conservation in PEAK3 orthologues (marked *) that may form a binding site for a PEAK3/Grb2 interactor.

We next sought to obtain structural data to better understand the mode of interaction of PEAK3 with Grb2. To do this we utilized a synthetic 7-mer pY phospho-peptide encompassing this SH2 motif (TpYSNLGQ; corresponding to PEAK3^23-29^-pY24 and PEAK1^1106-1112^-pY1107), hereafter ‘SH2-pY peptide’. We have previously shown by isothermal titration calorimetry (ITC) that this SH2-pY peptide binds to both full-length Grb2 (Grb2^FL^, *K*_D_ 2.6 µM) and CrkII (CrkII^FL^, *K*_D_ 7.8 µM)^14^.

Using an approach developed for apo-Grb2^FL^ ^38^, we succeeded in crystallizing a complex of human Grb2^FL^:PEAK SH2-pY peptide and solved an X-ray structure to a resolution of 2.7 Å (Table 1 and Fig. 2b-d). In our structure, Grb2^FL^ is present as a dimer as seen in the structure of apo-Grb2^FL^ (Protein Data Bank (PDB) ID: 1GRI)^38^, with the SH2-pY peptide bound to the SH2 domain of only one copy of Grb2 (chain B) (Fig. 2). Close crystal packing within the asymmetric unit leaves only this SH2 site (chain B) available for phosphopeptide binding (Fig. 2b-c). Our structure confirms that the PEAK3/PEAK1 SH2-pY peptide adopts a characteristic β-turn, with the expected recognition of the phosphotyrosine moiety and the specificity-determining (+2) N residue forming three important hydrogen bonds to the Grb2 backbone (R86, H107 and L109), consistent with other Grb2^SH2^:**pY**X**N**X peptide structures (eg. PDB ID: 1TZE)^39^. Whilst no density was apparent for the C-terminal (-GQ) residues of the peptide, we noticed a potential surface groove directly adjacent on Grb2^SH2^. In our orthologue multiple sequence alignment (MSA) data there is an extended region of high sequence conservation in PEAK3 and PEAK1 C-terminal to this SH2 motif (Fig. 2e). This corresponded to a predicted helical structure in AlphaFold2 (AF2)^40,41^ models of PEAK3 and PEAK1 (Extended Data Fig. 2a-b). Based on this, we spliced the peptide structure from our Grb2^FL^: PEAK SH2-pY peptide crystal structure with the AF2 model coordinates of the adjacent helical region (PEAK3^28-38^) to form an extended pY peptide model (PEAK3^23-38^-pY24) and undertook docking of this spliced peptide model into Grb2^FL^ using HADDOCK^42^. The resulting model of Grb2^FL^:PEAK3^23-38^-pY24 (Fig. 2e) reveals a compelling extended helix/groove interaction of the longer PEAK3 phosphopeptide with Grb2^FL^, with significant shape complementarity. Interestingly, whilst some of the highly conserved residues on PEAK3 are buried in the Grb2^SH2^ interface, others project into solvent presenting a new interaction surface. We speculate this creates a new conserved surface that might be involved in recruitment of an additional interactor to the Grb2:PEAK3-pY24 complex (and similarly for PEAK1) and may potentially also enhance selectivity for Grb2 over CrkII^SH2^.

**Table 1.**
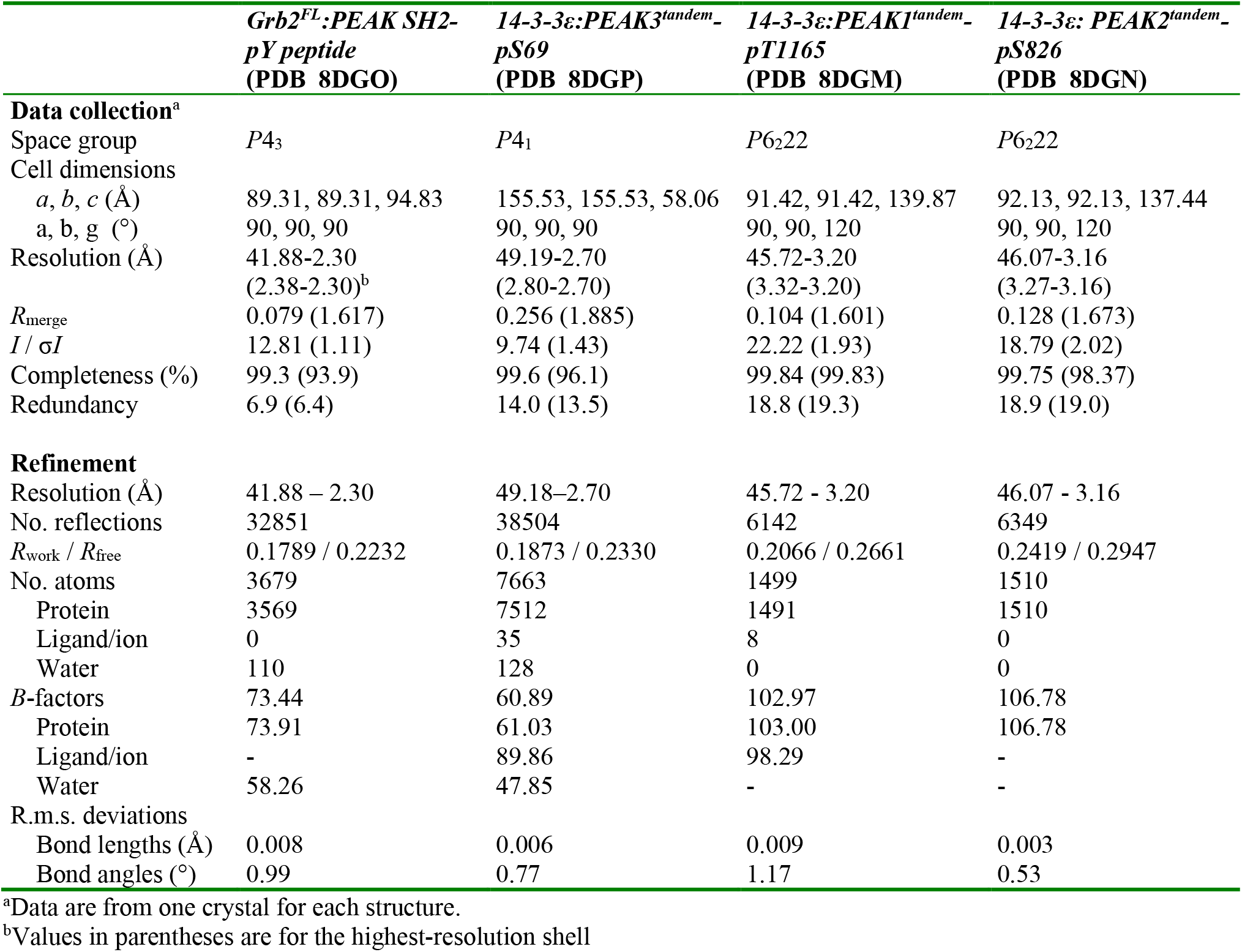
Data collection and refinement statistics.

### PEAK proteins bind CrkII^NSH3^ with similar affinity (low micromolar range)

Having made significant inroads to structurally and biophysically elucidate Grb2 binding at the PEAK3 SH2 site, we next turned our attention to the conserved ‘tandem site’ we identified on PEAKs, containing a CrkII^NSH3^ PRM and putative 14-3-3 binding site. Cellular studies show qualitatively that all PEAKs can interact with CrkII, requiring: (i) the conserved PRM within the ‘tandem site’ (matching the consensus for CrkII^NSH3^; PxLPxK); as well as (ii) SHED-mediated PEAK dimerization^4,14,32^. However, PEAK family members have non-identical sequence at this tandem site and to our knowledge no study has yet assessed the relative affinity of each for the CrkII^NSH3^, which is important to contextualize interaction hierarchy.

We therefore addressed this by generating a series of synthetic peptides that encompass the PRM from the tandem site of each PEAK protein - PEAK3 (residues 54-66), PEAK1 (residues 1150-1162), PEAK2 (residues 809-821) – as well as another PRM in PEAK2 similar to the consensus sequence, PEAK2 (residues 709-721), and determined their affinity towards the immobilized CrkII^NSH3^ domain by Surface Plasmon Resonance (SPR) (Fig. 3a and Extended Data Fig. 3a, Supplementary Information). We confirm that PEAK1^1150-1162^, PEAK2^809-821^ and PEAK3^54-66^ peptides all bind CrkII^NSH3^ with comparable affinity and fast on/off kinetics (measured dissociation constant, *K*_D_ 0.9 – 2.4 µM). Whilst the affinity of PRM/SH3 interactions can vary (1-100 µM range)^43^, this is comparable to CrkII^NSH3^ affinity for other PRMs that also closely match the CrkII^NSH3^ consensus sequence, such as the Abl^758^ PRM (reported *K*_D_ 1.7 µM).^44^ We also show that PEAK2^709-721^ represents a potential second, lower affinity binding site for CrkII^NSH3^ on PEAK2 (*K*_*D*_: 13 µM) (Fig. 3a and Supplementary Information). We observed no binding to CrkII^CSH3^ (data not shown). Comparison of the PEAK3^54-66^ peptide sequence to the crystal structure of the Abl PRM^758^ with CrkII^NSH3^ (PDB: 5IH2)^44^ (Fig. 3b) highlights the role that the Lys62 residue (-3 position) and the complementary electrostatic potential play in determining CrkII^NSH3^ specificity, as previously described for Abl^44,45^.

**Fig. 3:**
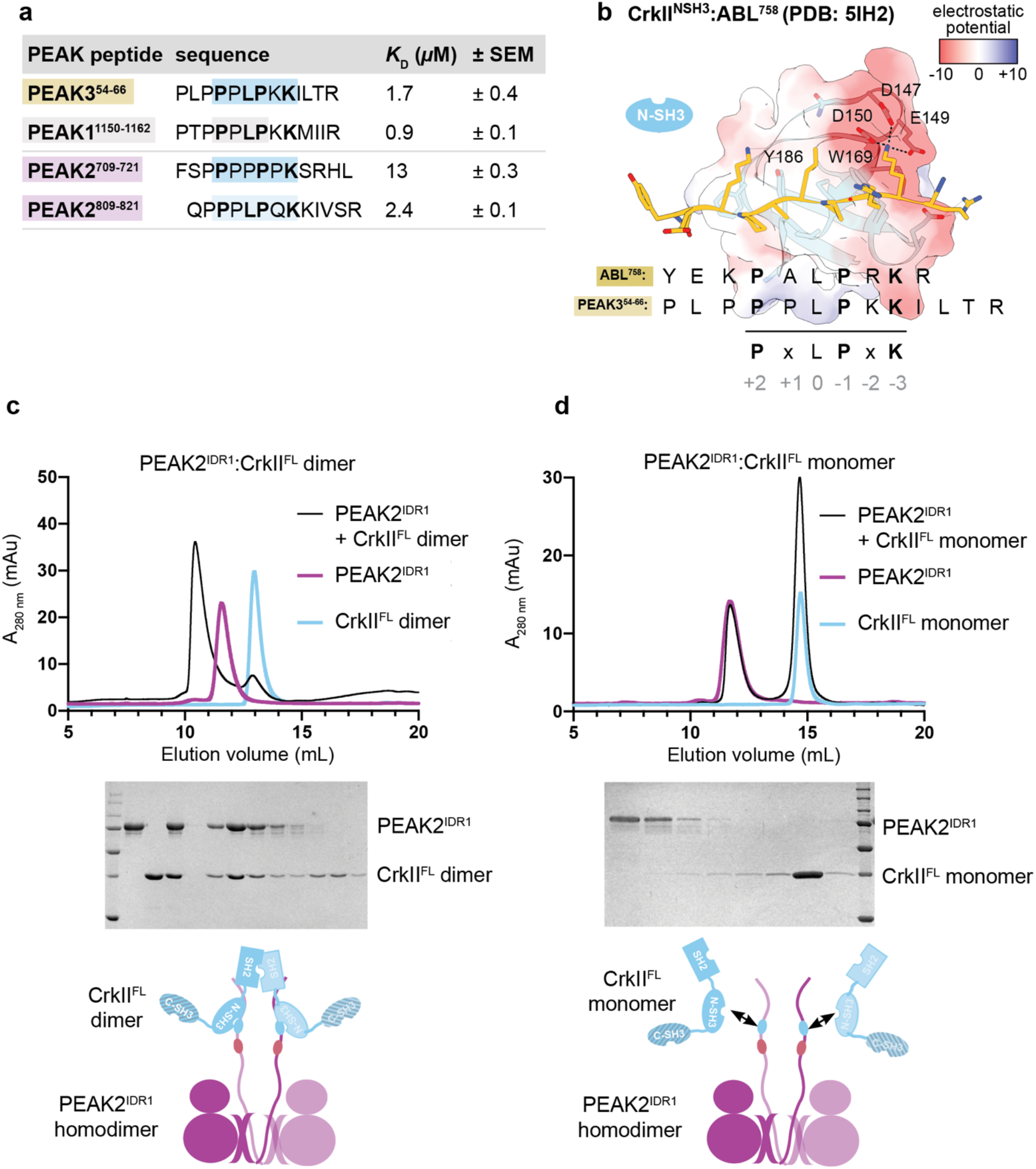
Structural and biophysical analysis of PEAK/CrkII^NSH3^ interaction. **(a)** SPR data summarizing measured steady-state binding affinity (*K*_D_) of synthetic PEAK PRM peptides towards immobilized CrkII^NSH3^ domain. **(b)** Depiction of the published CrkII^NSH3^:Abl^758^ structure (PDB: 5IH2)^44^, showing key residues within the CrkII^NSH3^ consensus motif in Abl^758^ (peptide shown in gold stick representation) crucial for high affinity binding to CrkII^NSH3^ (main chain light blue; CrkII^NSH3^ surface is coloured by electrostatic surface potential calculated using UCSF Chimera v 1.16; blue = positive, white = hydrophobic, red = negative). Aligned is the sequence of PEAK3^54-66^, showing key conserved residues of the CrkII^NSH3^ motif. **(c, d)** Interaction studies with recombinant PEAK1^IDR1^/PEAK2^IDR1^ and CrkII^FL^ underscore the role of avidity for high affinity binding of CrkII^FL^ to PEAK dimers; (c) Incubation of PEAK2^IDR1^ with CrkII^FL^ dimer at 1:1.3 ratio results in a complex formation while (d) incubation with CrkII^FL^ monomer does not result in a complex, as confirmed by SDS-PAGE analysis of SEC eluted fractions. CrkII^FL^ dimer and monomer in light blue; PEAK2^IDR1^ in pink; complex of PEAK2^IDR1^ dimer:CrkII^FL^ dimer in black line (see Extended Data Fig. 3d for PEAK1^IDR1^ dimer:CrkII^FL^ dimer complex).

### Avidity enables stable PEAK interactions with CrkII

We next looked to examine the role of dimerization in CrkII binding to PEAKs. As our initial SPR studies had utilized isolated peptides and only the CrkII^NSH3^ domain, we shifted to utilize recombinant CrkII^FL^ and dimeric PEAK1, PEAK2 and PEAK3. CrkII^FL^ was expressed and purified from *E*.*coli* as described previously^14^. Interestingly, whilst the majority of CrkII^FL^ is monomeric, approximately 10% of the total isolated protein purifies as a stable dimer (Extended Data Fig. 3b-c). Dimerization has been described for other adapters such as Grb2^46^ and the related protein CrkL (mediated by the C-terminal SH3 domain^47^), however to our knowledge this has not been reported to date for CrkII. We confirm that, in contrast to CrkL, the dimerization of CrkII^FL^ we observe is mediated by the SH2 domain, as recombinant C-terminally truncated constructs CrkII^ΔCSH3^ (SH2-NSH3) and CrkII^SH2^ each form a similar proportion of dimer (Extended Data Fig. 3b-c). As MS analysis of both monomeric and dimeric CrkII^FL^ shows no PTMs (See Source Data) we believe the dimer we observed to be most likely due to a domain-swap within the SH2 domain.

We next focused on PEAK1 and PEAK2, as the SHED-PsK domains of these have previously been purified and crystallised^18,37^ and designed longer constructs, PEAK1^IDR1^ and PEAK2^IDR1^, harboring the ‘tandem motif’^37^. We recently reported the expression/purification of these longer constructs from insect cells^37^. Consistent with our previous findings for PEAK1/2^18^, PEAK1^IDR1^ and PEAK2^IDR1^ each purify as a homodimer (Fig. 3c-d and Extended Data Fig. 3d). Purified PEAK1^IDR1^ and PEAK2^IDR1^ were then analyzed for their ability to interact with either monomeric or dimeric CrkII^FL^ on SEC followed by SDS-PAGE (Fig. 3c-d, Extended Data Fig. 2). Strikingly, whilst the PEAK2^IDR1^ dimer:CrkII^FL^ monomer complex eluted as two peaks at the expected elution volume for each component, consistent with the anticipated low-affinity complex (micromolar range with fast on/off kinetics), PEAK2^IDR1^ dimer:CrkII^FL^ dimer complex showed a marked peak shift to an earlier elution volume, indicative of a stable interaction (Fig. 3c). SDS-PAGE confirmed that this complex contained both PEAK2^IDR1^ and CrkII^FL^ dimers (Fig. 3c). Similar results were obtained for PEAK1^IDR1^ dimer:CrkII^FL^ dimer (Extended Data Fig. 3d). SEC coupled to Multi-Angle Light Scattering (SEC-MALS) analysis of the purified PEAK2^IDR1^ dimer:CrkII^FL^ dimer complex estimated a mass of 192 kDa, close to the expected mass of 198 kDa for a stoichiometric (2:2) PEAK2^IDR1^ dimer:CrkII^FL^ dimer complex (Extended Data Fig. 3e).

The marked differences in complex stability observed between binding of dimeric and monomeric CrkII^FL^ to dimeric PEAK1 and PEAK2 reveal an important role for avidity for stable recruitment to PEAKs, helping to rationalize the dimerization requirement observed in cellular studies^48^.

### PEAK3 can bind 14-3-3 via the tandem site 14-3-3 motif (pS69)

We next wanted to extend these studies to recombinant PEAK3^FL^. Interestingly, PEAK3^FL^ from insect cells was purified as a stable heteromeric complex with two additional interactors, identified by MS as insect cell derived 14-3-3 proteins (14-3-3ε,ζ heterodimer) (Fig. 4a and Source Data)^37^. This was consistent with our SLiM identification of a highly conserved putative 14-3-3 motif present in the PEAK3 IDR. Co-purification of 14-3-3 from insect cells has been observed for other kinases with 14-3-3 binding sites (e.g. BRAF, LRKK2)^29,49-51^. Proteomic analysis of the purified PEAK3^FL^:14-3-3ε,ζ complex confirmed that recombinant PEAK3 purified from insect cells was phosphorylated only at the 14-3-3 motif (pS69) within the tandem site (Extended Data Fig 4a). Despite apparent conservation of this tandem 14-3-3 motif across the PEAK family, we observed differences in both phosphorylation and 14-3-3 interaction at this motif between PEAK family members. Proteomic analysis of insect cell derived PEAK2^IDR1^ showed phosphorylation of the tandem site 14-3-3 motif (RAA<u>S</u>SP), whereas phosphorylation at this motif was not observed for PEAK1^IDR1^ (RAN<u>T</u>EP), moreover co-purification with 14-3-3 was only observed for PEAK3^FL^ (Extended Data Fig. 4a). To complement these data and further characterize the contribution of PEAK3 pS69 in recruiting 14-3-3 in cells, we performed anti-HA IPs from HEK293 cells expressing HA-tagged PEAK3^FL^ WT (HA/PEAK3^FL^) and PEAK3^FL^-S69A mutant (HA/PEAK3^FL^-S69A; Fig. 4b and Extended Data Fig. 4b). These experiments confirmed that while HA-tagged PEAK3^FL^ WT can co-immunoprecipitate endogenous 14-3-3, this interaction was abolished with HA-tagged PEAK3 ^FL^-S69A, indicating that S69 is the primary cellular 14-3-3 interaction site for PEAK3.

**Fig. 4:**
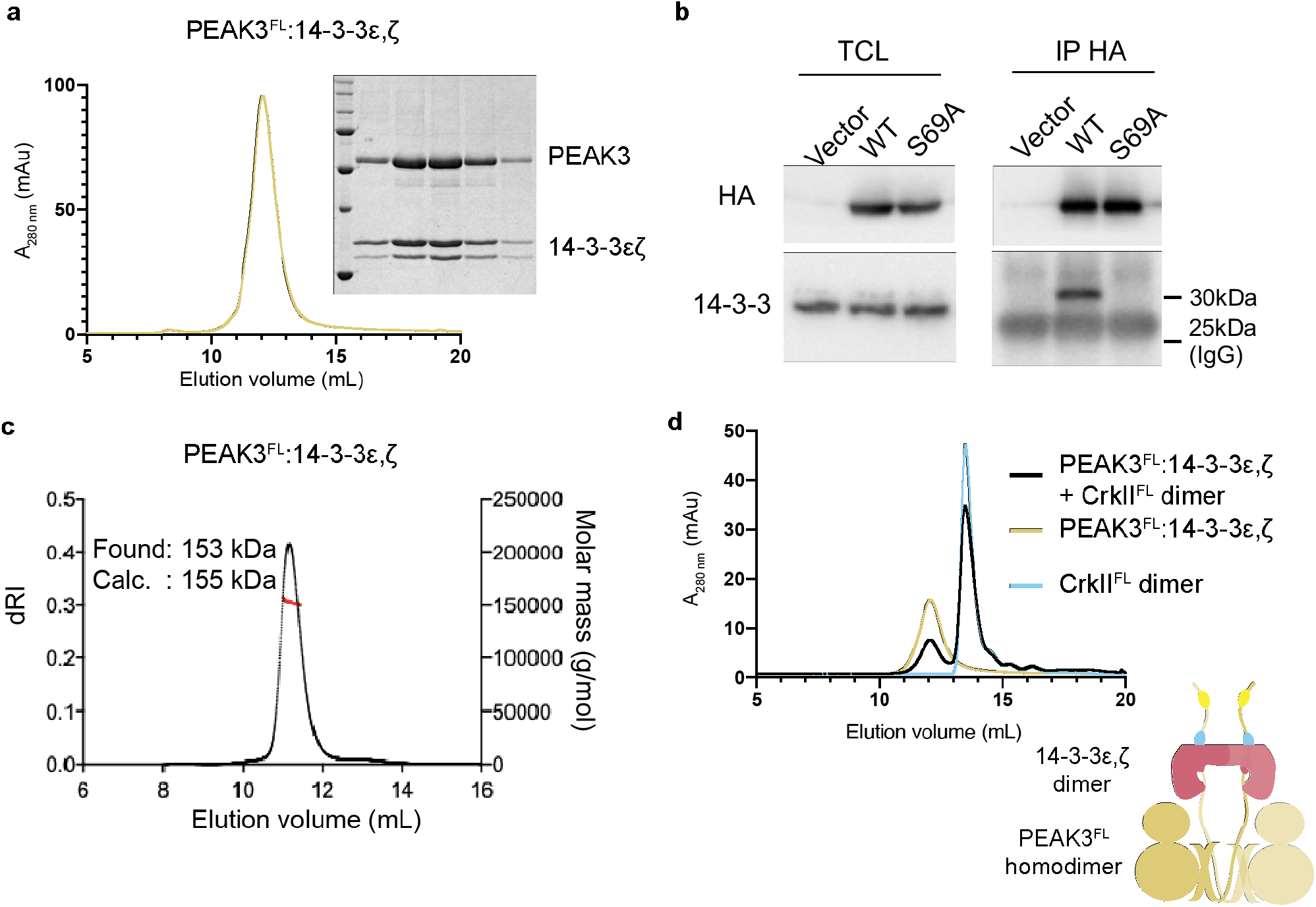
PEAK3:14-3-3 forms a stable high affinity heterocomplex. **a**, Recombinant PEAK3^FL^ from insect cells elutes on Size Exclusion Chromatography (SEC; S200 10/300) as a high affinity stoichiometric complex of dimeric PEAK3 with a 14-3-3ε,ζ heterodimer, results supported by SDS-PAGE analysis of eluted fractions (See Extended Data Fig. 4a, Source Data). **b**, Immunoprecipitation of PEAK3 WT and PEAK3 S69A showing S69 is required for 14-3-3 co-immunoprecipitation from cells (n=3, see Extended Data Fig. 4a for biological repeats). **c**, SEC-MALS analysis of PEAK3^FL^-14-3-3ε,ζ heterodimer complex confirming the experimentally determined mass closely matches the mass of 155 kDa expected for a stoichiometric dimer:dimer complex. **c**, SEC profile (S200 10/300) of recombinant PEAK3^FL^-14-3-3ε,ζ heterodimer pre-incubated with CrkII^FL^ dimer in a 1:1.2 molar ratio) showing no complex formation.

### PEAK3:14-3-3 form a stable dimer:dimer heterocomplex

We recently showed that PEAK3 can form homodimers in a cellular context^14^. To study the effect of PEAK3 dimerization on 14-3-3 binding in cells, we co-expressed in HEK293 cells Flag-tagged PEAK3^FL^ WT with HA-tagged versions of PEAK3^FL^ WT or PEAK3^FL^ dimerization mutants (L146E, A436E, C453E)^14,18^. Immunoprecipitation and western blotting experiments confirmed that the inability of these mutants to undergo homodimerization is associated with a lack of recruitment of 14-3-3 (Extended Data Fig. 4c), demonstrating that 14-3-3 association to PEAK3 is also dependent on PEAK3 dimerization. These findings build on our previous data^14^ (and reports from the Jura laboratory^4^) demonstrating the dimerization dependent recruitment of other interactors such as CrkII and ASAP1 to PEAK3.

We next used SEC-MALS to confirm the stoichiometry of our recombinant PEAK3^FL^:14-3-3ε,ζ heteromeric complex, as homo or heterodimers of 14-3-3 will contain two separate phosphopeptide binding sites. A single symmetrical peak with an experimental mass of 153 kDa was observed, suggesting a stable PEAK3^FL^:14-3-3ε,ζ dimer:dimer complex (expected mass 155 kDa) (Fig. 4c). We next performed SEC interaction studies of the PEAK3^FL^:14-3-3ε,ζ complex with the CrkII^FL^ dimer. Interestingly, while CrkII^FL^ complexes with PEAK1^IDR1^ or PEAK2^IDR1^ (Fig. 3c-d), the PEAK3^FL^:14-3-3ε,ζ complex showed no complex formation with CrkII^FL^, despite the presence of the CrkII^NSH3^ binding motif in the PEAK3^FL^ construct (Fig. 4d).

### PEAK3 shows a high affinity phosphodependent interaction with 14-3-3 at the tandem site

Given the proximity of the 14-3-3 motif and CrkII^NSH3^ motifs within the tandem site, we next wanted to compare binding affinity of 14-3-3 and CrkII ^NSH3^ at each respective site (Fig. 5 and Extended Data Fig. 5). We generated peptides of PEAK3, PEAK1 or PEAK2 that encompass the tandem site (CrkII^NSH3^ and 14-3-3 motif), phosphorylated or non-phosphorylated at the 14-3-3 motif and conducted biophysical interaction studies by SPR with purified 14-3-3 and CrkII proteins (Extended Data Fig. 5a and Supplementary Information). We selected, as an initial subset, four human 14-3-3 isoforms (ɣ, ε, η, σ) out of the 7 human isoforms, as those were the most highly represented in proteomic interaction analysis with PEAK proteins in cells^4,20^. SPR analysis of PEAK3 tandem peptides confirmed that phosphorylated PEAK3^tandem^-pS69 peptide binds with high affinity to immobilized 14-3-3ɣ (SPR dissociation constant, *K*_D_ = 0.19 µM), whereas the corresponding non-phosphorylated PEAK3^tandem^-S69 peptide shows negligible binding (*K*_D_ > 10 µM) (Fig. 5a).

**Fig. 5:**
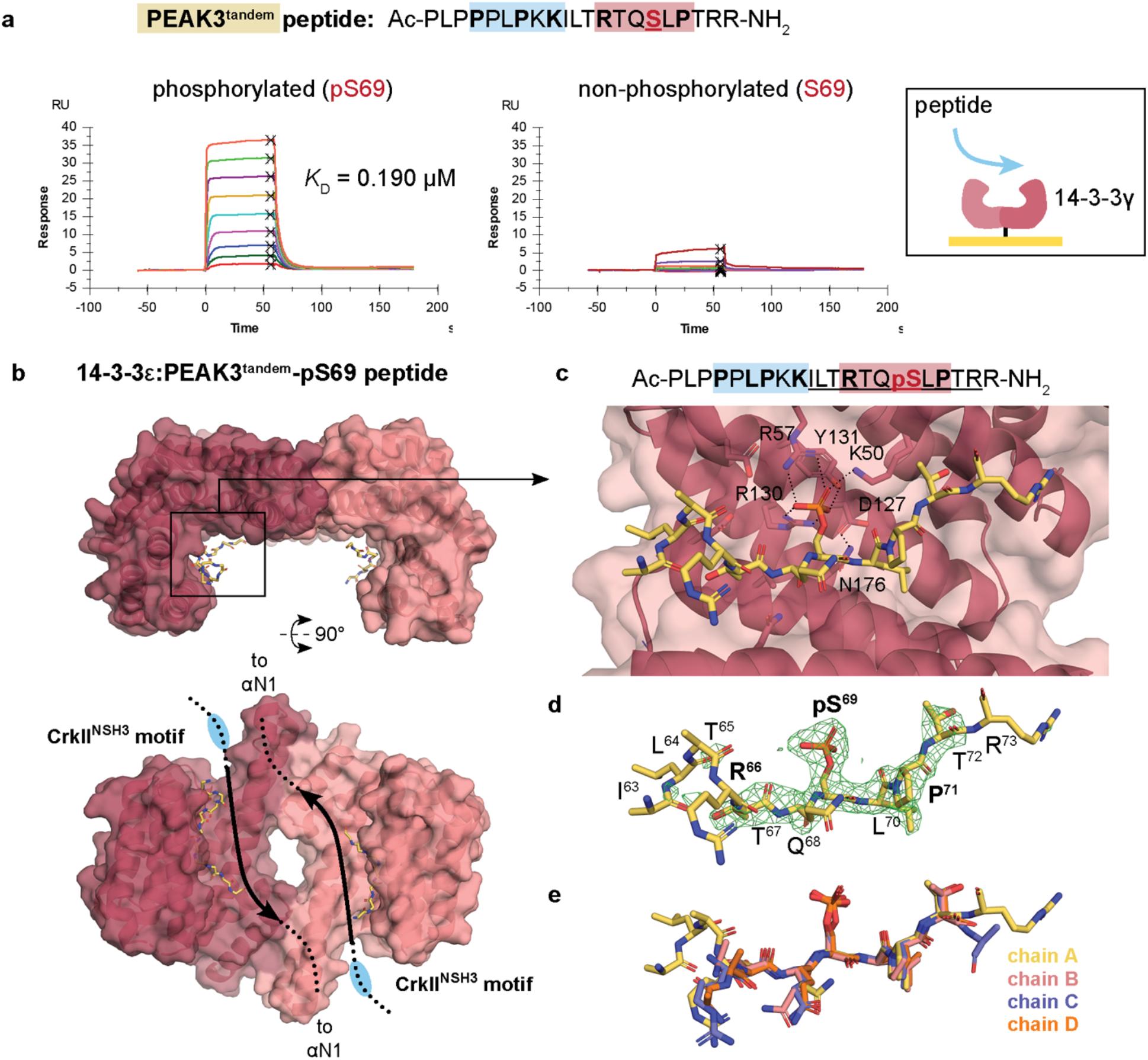
Structural and biophysical analysis of PEAK3/14-3-3 interaction. **a**, Binding of PEAK3^tandem^-pS69 (left) and PEAK3^tandem^-S69 peptides (right) to 14-3-3ɣ as measured by SPR. **b**, Overall structure of 14-3-3ε: PEAK3^tandem^-pS69 peptide showing 14-3-3 dimer antiparallel arrangement with each monomer (pink cartoon/surface) bound to a single copy of the PEAK3^tandem^-pS69 peptide (yellow sticks). Chain A of 14-3-3ε is in dark pink and chain B is in light pink. **c**, Zoom in highlighting peptide groove and showing the central pS69 residue of the PEAK3 phosphopeptide interacting with K50, R57, Y131 and R130. **d**, Unbiased Fo-Fc omit map (green mesh, contoured at 3.0 σ) showing peptide density prior to modelling and final modelled PEAK3^tandem-^pS69 phosphopeptide (yellow sticks). **e**, Superposition of the peptide modelled in each 14-3-3 monomer (chains A-D, sticks) in the asymmetric unit.

We next compared SPR binding of PEAK3, PEAK1 and PEAK2 tandem peptides to human 14-3-3ɣ, 14-3-3ε, 14-3-3η and 14-3-3σ. For each isoform, phosphodependent binding was observed and the PEAK3^tandem^-pS69 phosphopeptide consistently showed approximately 10 fold tighter affinity (*K*_D_ ∼ 0.16-0.78 µM) than those of PEAK1^tandem^-pT1165 or PEAK2^tandem^-pS826 (*K*_D_ ∼ 1-10 µM) (Extended Data Fig. 5c). This difference was also apparent in the comparatively slower dissociation kinetics for 14-3-3:PEAK3^tandem^-pS69 complexes relative to PEAK1 or PEAK2 tandem peptides (Extended Data Fig. 5c and Supplementary Information).

To gain molecular details of the PEAK3 interaction with 14-3-3, we determined the crystal structure of a 14-3-3ε:PEAK3^tandem^-pS69 complex to 2.5 Å resolution (Table 1). This complex crystallized with two 14-3-3ε dimers in the asymmetric unit (ASU), each bound to a single copy of the PEAK3^tandem^-pS69 peptide. In each symmetric 14-3-3ε dimer, the PEAK3^tandem^-pS69 peptide is present in the peptide binding groove in an overall antiparallel arrangement (Fig. 5b), with clear unbiased electron density for the central phosphoserine residue (PEAK3 pS69) recognized by 14-3-3ε as well as adjacent residues of the PEAK3 motif (approximately 7 residues in total). Notably, interactions typical of such 14-3-3 complexes are observed: the PEAK3 pS69 phosphate group is coordinated by 14-3-3ε residues K50, R57, R130 and Y131 and 14-3-3ε D127 and N176 interact to enable N176 to form a hydrogen bond to the backbone amide -NH of PEAK3 L70. Each copy of the PEAK3^tandem^-pS69 peptide within the ASU adopts a similar conformation (Fig. 5c-e, Extended Data Fig. 5b and Table 1). We also solved two crystal structures of 14-3-3ε: PEAK1^tandem^-pT1165 and 14-3-3ε:PEAK2^tandem^-pS826 complexes, which were lower resolution (3.1Å) but similarly confirmed phosphorecognition and a canonical mode of binding. In each of the three 14-3-3:tandem peptide structures, no clear electron density was observed for the adjacent CrkII^NSH3^ PRM likely due to flexibility and crystallographic averaging.

### The 14-3-3 site on PEAK3 represents a molecular switch for CrkII binding

To biophysically evaluate potential positive or negative cooperativity between these two adjacent binding sites in the PEAK3 tandem motif, we turned to sequential binding studies by ITC (Fig. 6a-b and Extended Data Fig. 6a-d). In these experiments, the PEAK3 tandem peptide (either phosphorylated or non-phosphorylated) was loaded in the cell and protein (either 14-3-3ɣ or CrkII^NSH3^) in the syringe. First a titration of 14-3-3ɣ was measured, to saturate the first site on the peptide (binary experiment). The tandem peptide/14-3-3ɣ complex was retained in the cell and a second titration measured for CrkII^NSH3^ in the presence of 14-3-3ɣ (ternary experiment) (Fig. 6). Corresponding sequential titrations were also conducted using buffer (first injection) followed by CrkII^NSH3^ (second injection) (binary experiment). Direct SPR binding studies for each PEAK3^tandem^ peptide to immobilized CrkII^NSH3^ were also undertaken. The results for ITC and SPR binding experiments for the PEAK3^tandem^ peptides are summarized in Table 2.

**Table 2:**
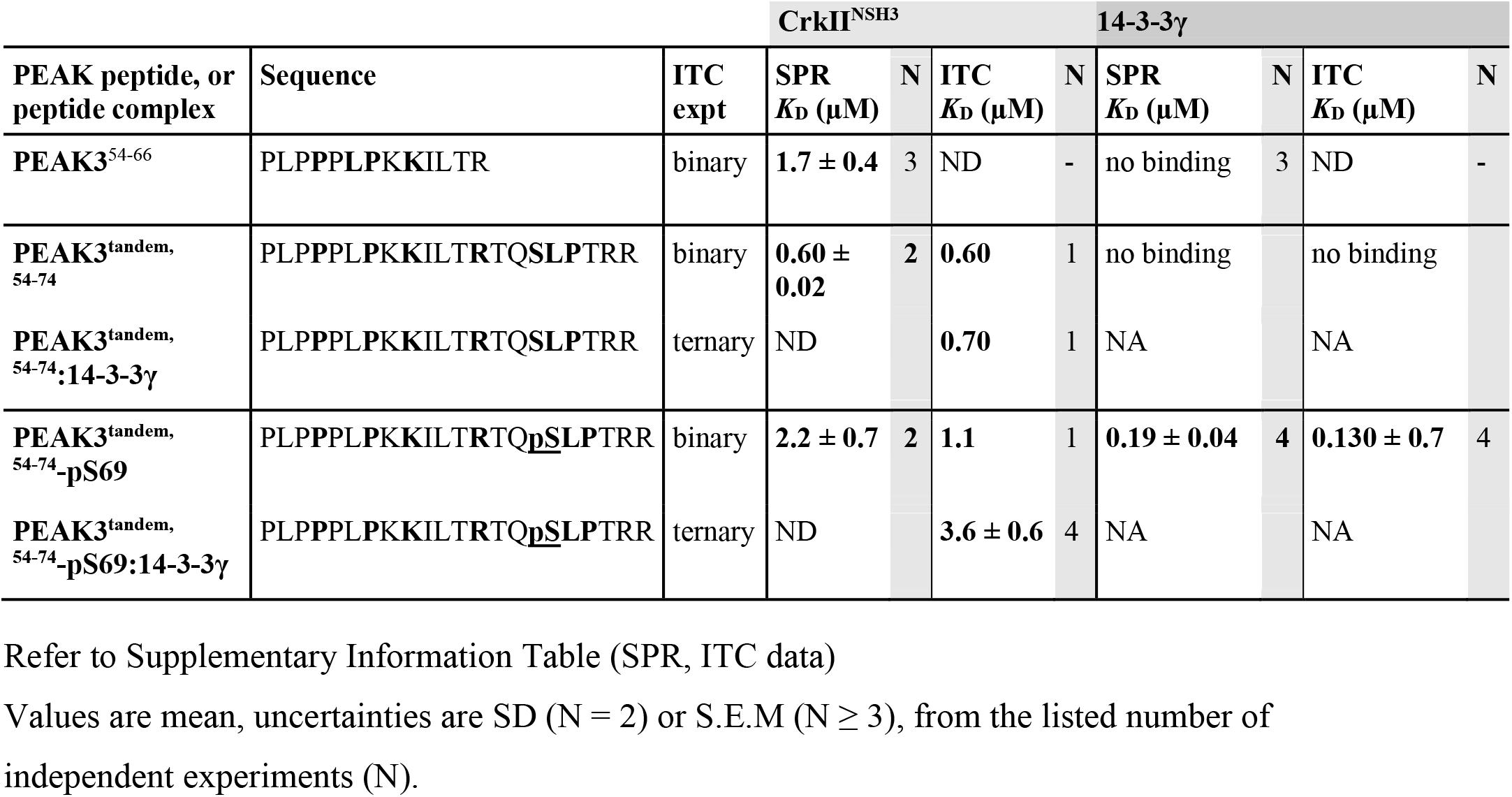
Summary of ITC sequential binding studies data.

| PEAK peptide, or peptide complex | Sequence | ITC expt | CrkII<sup>NSH3</sup> | | | | 14-3-3γ | | | |
| --- | --- | --- | --- | --- | --- | --- | --- | --- | --- | --- |
| | | | SPR $K_D$ (μM) | N | ITC $K_D$ (μM) | N | SPR $K_D$ (μM) | N | ITC $K_D$ (μM) | N |
| PEAK3<sup>54-66</sup> | PLPPPLPKKILTR | binary | 1.7 ± 0.4 | 3 | ND | - | no binding | 3 | ND | - |
| PEAK3<sup>tandem, 54-74</sup> | PLPPPLPKKILTRTQSLPTRR | binary | 0.60 ± 0.02 | 2 | 0.60 | 1 | no binding | | no binding | |
| PEAK3<sup>tandem, 54-74</sup>:14-3-3γ | PLPPPLPKKILTRTQSLPTRR | ternary | ND | | 0.70 | 1 | NA | | NA | |
| PEAK3<sup>tandem, 54-74</sup>-pS69 | PLPPPLPKKILTRTQ<u>p</u>SLPTRR | binary | 2.2 ± 0.7 | 2 | 1.1 | 1 | 0.19 ± 0.04 | 4 | 0.130 ± 0.7 | 4 |
| PEAK3<sup>tandem, 54-74</sup>-pS69:14-3-3γ | PLPPPLPKKILTRTQ<u>p</u>SLPTRR | ternary | ND | | 3.6 ± 0.6 | 4 | NA | | NA | |

Refer to Supplementary Information Table (SPR, ITC data)
Values are mean, uncertainties are SD (N = 2) or S.E.M (N ≥ 3), from the listed number of independent experiments (N).

**Fig. 6:**
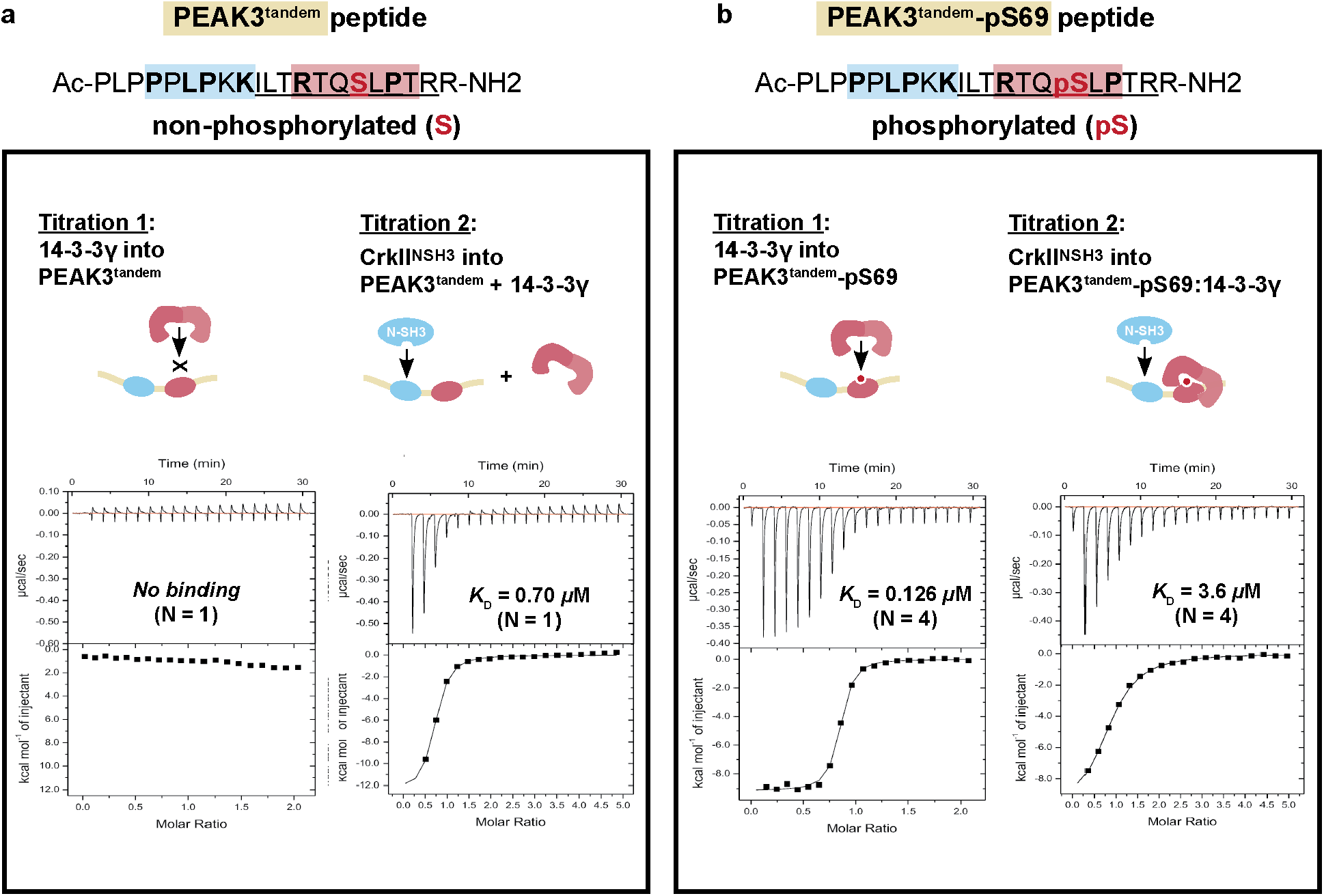
14-3-3ɣ binding to PEAK3^tandem^ peptide reduces the affinity of CrkII^NSH3^ binding to the adjacent CrkII motif. **a, b**, Sequential ITC binding studies using PEAK3^tandem^-S69 non-phosphorylated (a) and PEAK3^tandem^-pS69 phosphorylated (b) peptides, purified 14-3-3ɣ and purified CrkII^NSH3^ domain. High affinity binding of 14-3-3 ɣ to PEAK3^tandem^-pS69 peptide (b) reduces the affinity of CrkII^NSH3^ binding to the adjacent CrkII motif at the tandem site (negative cooperativity). See Extended Data Fig. 6a-d.

For the non-phosphorylated PEAK3^tandem^ peptide, both SPR and ITC yield an affinity (*K*_D_) of 0.60 µM for binding to CrkII^NSH3^, whilst no appreciable binding of this peptide to 14-3-3ɣ is observed. Similarly, the affinity of this peptide for CrkII^NSH3^ was unaffected by first pre-saturating with 14-3-3ɣ (ITC *K*_D_ 0.70 µM). In contrast, phosphorylated PEAK3^tandem^-pS69 peptide showed strong binary binding to 14-3-3ɣ by either method (SPR *K*_D_ 0.19 µM, ITC *K*_D_ 0.13 µM), but modestly weaker binary binding to CrkII^NSH3^ relative to the non-phosphorylated peptide (SPR *K*_D_ 2.2 µM, ITC *K*_D_ 1.1 µM). It is possible that the phosphate group reduces electrostatic complementarity to the adjacent negatively charged patch on the CrkII^NSH3^ binding cleft^52^. When the phosphorylated PEAK3^tandem^-pS69 peptide was first pre-complexed with 14-3-3ɣ, binding of CrkII^NSH3^ was further weakened (ITC *K*_D_ 3.6 µM). This represents a 6-fold loss in affinity for the PEAK3 tandem peptide towards CrkII^NSH3^ following pS69 phosphorylation and 14-3-3 binding, demonstrating negative cooperativity in these interactions at the tandem site. In the context of a dimeric scaffold (dimer:dimer of PEAK3:14-3-3ε,ζ) it is expected this negative cooperativity will be further pronounced due to avidity effects, which supports the inability of the CrkII^FL^ dimer to show measurable complex formation with PEAK3:14-3-3ε,ζ *via* SEC.

Consistent with these biochemical and biophysical data, in our HEK293 cellular immunoprecipitation experiments in using wildtype HA-tagged PEAK3^FL^ and PEAK3^FL^-S69A mutant, we observed that alongside the complete loss of 14-3-3 association for the PEAK3^FL^-S69A mutant was a modest but significant increase in level of associated CrkII, relative to PEAK3^FL^-WT (Extended Data Fig. 4b). Importantly, this is despite the level of CrkII associated with PEAK3^FL^-WT (or S69A) in these experiments likely being a composite of a number of multivalent CrkII interactions with multiple partners, including potentially interaction of CrkII with heterodimeric PEAK3/PEAK1 or PEAK3/PEAK2 via the CrkII^NSH3^ site, PEAK3-pY24 motif via the CrkII^SH2^ site and possible bridging *via* other adapters such as Grb2, as we have shown previously^14^.

### Building a model for PEAK3/CrkII regulation via the tandem site 14-3-3 motif

Taken together, our data provide the molecular basis for a model of PEAK3 regulation, in which the ‘tandem site’ we identified within the relatively short IDR of PEAK3 acts a molecular switch for CrkII binding (Fig. 7a), acting via 14-3-3. PEAK3, PEAK1 and PEAK2 (as homo- or heterodimers) can all bind CrkII at the conserved tandem site (*K*_D_ ∼ 1-2 µM for each 1:1 interaction with CrkII^NSH3^ domain, but higher effective *K*_D_ of PEAK dimer for clustered CrkII to avidity) (Fig. 7a(i)). Phosphorylation at PEAK3 S69 (tandem site) generates a high affinity binding site for dimeric 14-3-3 proteins (*K*_D_ ∼ 0.1-1 µM for at each site, further enhanced by avidity due to dimer/dimer interaction). This enables formation of a highly stable PEAK3:14-3-3 dimer:dimer complex (Fig. 7a(ii),b). Due to negative cooperativity between 14-3-3 and CrkII^NSH3^ binding at each tandem site, the PEAK3 interaction with CrkII is destabilized (Fig. 7a(iii)). 14-3-3 thus appears to be a key regulator of PEAK3 activity and modulates signalling via CrkII, with the crucial switching event being phosphorylation at the PEAK3 tandem site S69 motif. As an intact PEAK3 binding site for CrkII^NSH3^ is also shown to be crucial for Src/SFK-mediated phosphorylation of PEAK3 and binding of Grb2/ASAP1 at the adjacent Y^24^ site, this may also implicate 14-3-3 binding in regulation of this process (Fig. 7a(iv)).

**Fig. 7:**
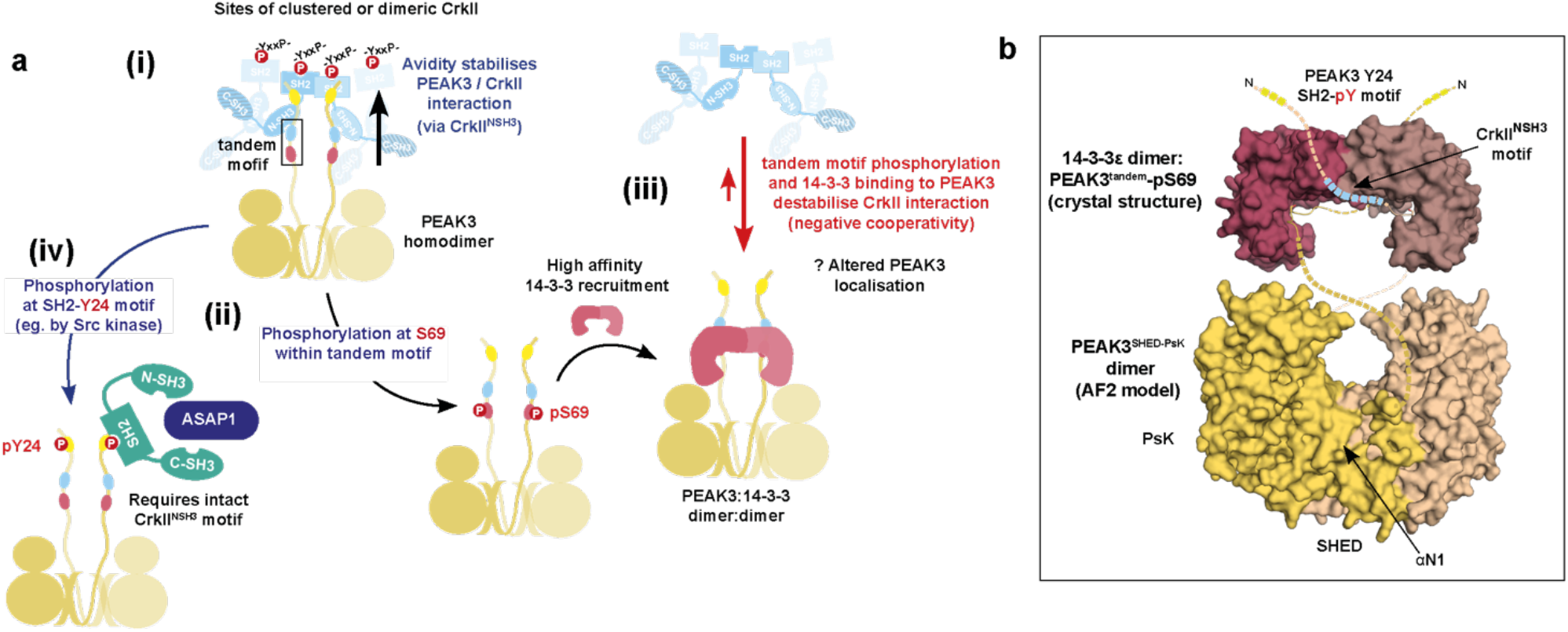
Proposed model for PEAK3/Grb2 and PEAK3/CrkII regulation via the tandem site 14-3-3 motif. **a, (i)** Avidity stabilizes PEAK3/CrkII interaction with clustered CrkII (interacting via the CrkII^NSH3^ and tandem motif PRM of the PEAK3 dimer). CrkII is likely to be clustered by interaction with other adapters/scaffolds, such as SH2 domain-mediated binding to partner proteins containing multiple adjacent pYxxP motifs (eg. p130cas/NEDD9, paxillin), but possibly also via dimerization (eg. SH2-mediated as we identify for recombinant protein) **(ii)** Phosphorylation of PEAK3 at S69 creates a high-affinity binding site for 14-3-3, leading to the formation of a highly stable PEAK:14-3-3 dimer:dimer. **(iii)** Phosphorylation/binding of 14-3-3 to PEAK3 destabilizes CrkII binding in the adjacent tandem site (negative cooperativity), leading to disruption of PEAK3/CrkII complex. This may terminate PEAK3/CrkII signaling and/or alter PEAK3 subcellular localization. **(iv)** Phosphorylation of PEAK3 at Y24 (eg. by Src kinase) recruits Grb2 via its SH2 domain leading to association with ASAP1 and PYK2 and potential PYK2/ASAP1-mediated activation of the PI3K/AKT pathway. **b**, Illustrative representation of a potential dimeric human PEAK3^FL:^14-3-3ε complex to illustrate relative scale and key sites of interaction. Depicted is the 14-3-3ε:PEAK3^tandem-pS69^ crystal structure (PDB 8DGP) and a structural model of the PEAK3^SHED-PsK^ dimer prepared from AlphaFold2 multimer modelling of a human PEAK3^FL^ dimer^70,71^. Unmodelled IDR linker regions are depicted as dotted lines. The precise arrangement of the PEAK3^FL^ dimer relative to the 14-3-3 dimer as shown is arbitrary, although some constraint on relative positioning is expected to be imparted by the short length of the PEAK3 IDR and antiparallel arrangement of interaction with 14-3-3.

## DISCUSSION

Our studies illuminate several aspects of the dynamic regulation of PEAK scaffolding functions, particularly in the context of integrin- or EGFR-mediated signaling (Fig. 7, Extended Data Fig. 7). We apply a combination of analytical techniques to characterize PEAK3 interactions with Grb2, CrkII and 14-3-3 via its N-terminal IDR, and the role of phospho-regulation within the PEAK3 IDR. We demonstrate how dimerization-induced avidity effects and molecular crowding can each modulate SH3/PRM interactions, including how these mechanisms of interaction are conserved or differ across the PEAK family.

Firstly, we provide new molecular insights to contextualize the interaction of PEAK3 and PEAK1 with Grb2/ASAP1^14,23^. We confirm that active Src, but not Abl, phosphorylates the conserved TYSNL SH2-motif (PEAK3^Y24^, PEAK1^Y1107^), supporting previous cellular studies,^14^ and thereby provides a binding site on either PEAK3 or PEAK1 for Grb2^SH2^ or possibly CrkII^SH2^ (Fig. 2). We provide a high-resolution structure of Grb2^FL^ characterizing the binding of PEAK1/PEAK3 at this site and identify a conserved interface mediated by the PEAK3 peptide when bound to Grb2 that provides a possible docking site for a PEAK/Grb2 interactor. ASAP1 is a possible candidate, as PEAK3 pY24 phosphorylation is required for both ASAP1 and Grb2 interaction with PEAK3^14^ and the conserved solvent-exposed residues of PEAK3 predicted from the Grb2^FL^ docking model (K^38^, P^36^, L^35^, R^31^) also resemble the linear consensus motif for the ASAP1 SH3 domain (KPxR)^53^ (Fig. 2). The observation that PEAK3 dimerization is required for Y^24^ phosphorylation (and Grb2 association)^14^ is also consistent with a model in which clustered CrkII assists to colocalize dimeric PEAK3 with an active kinase (e.g. Src) for efficient Y^24^ phosphorylation. Grb2 is also thought to undergo dynamic switching between dimeric (inactive) and monomeric (active, Sos-binding) states; a feature reported to be pivotal for Grb2 signalling in cancer ^46^. Further studies are needed to examine whether PEAK dimers can impact this process also.

Secondly, we examine how NSH3/PRM interactions and avidity together mediate PEAK3 interaction with CrkII, a key player in FA turnover. Our work clearly places PEAKs as recruited scaffolds within this context at FAs (Fig. 7a,b), together with PYK2 and active Src, other important downstream mediators of integrin signaling^54,55^. We show that the conserved PRM in the PEAK IDR tandem site alone confers modest affinity for CrkII^NSH3^ (SPR *K*_D_ ∼ 1 µM with a rapid off-rate; Fig. 3), which is essentially equivalent for all PEAKs. However, we demonstrate using monomeric and dimeric forms of CrkII^FL^ that SHED-dependent homodimerization (and by implication also hetero-dimerization of PEAKs)^4,17-20^ enables enhanced stability of PEAK complexes with dimeric CrkII due to avidity (Fig. 3; Extended Fig. 7b). These findings further support and rationalize data from this work and other reports^4,14^ demonstrating that mutation of key residues of the PEAK3 dimer interface disrupts PEAK3/CrkII interactions in cells, which would impact avidity of these complexes rather than the PRM/SH3 interaction.

We therefore propose that high avidity binding between the CrkII^NSH3^ motif and the conserved tandem site PRM on PEAK dimers is a general feature of PEAKs that enhances localization to FAs or other sites of CrkII clustering (Fig. 7a; Extended Fig. 7)^32,56^. Whilst we identify an SH2-mediated dimeric form of CrkII^FL^ from overexpression in *E. coli*, further work will be required to confirm whether this exists in a cellular context. However, as multivalency is a general feature of both CrkII and CrkL signaling complexes, our work has broader implications. A notable example is cell adhesion -mediated CrkII/CrkL interactions with the Cas family of scaffolds (p130Cas or NEDD9), each of which harbor flexible central substrate domains with multiple repeats (15 repeats in p130Cas) of the YxxP motif; a consensus binding sequence for SH2- and PTB-containing proteins such as CrkL, Nck, and SHIP2. Integrin-mediated cell adhesion allows Cas to activate Src/SFKs, which processively phosphorylate the Cas substrate domain following mechanosensitive changes in its accessibility, ultimately enabling multivalent CrkII recruitment^57,58^. In this manner, CrkII cooperates with p130Cas/NEDD9 in dynamic integrin signaling at FAs^59^. Bound and clustered CrkII then recruits other adapters/effectors of other pathways to amplify and diversify the signal including Dock180/Rac (regulation of migration) and C3G/Rap1 (regulation of cellular morphology and adhesion)^60^. Likewise, AXL-mediated NEDD9 phosphorylation is reported to recruit CrkII and thereby PEAK1, leading to altered FA dynamics in breast cancer cells^32^. This is consistent with roles for p130Cas/NEDD9 in tumor progression through regulating cytoskeletal dynamics^61^. Thus, although not specifically examined in our work, our findings with CrkII likely also have implications for PEAK interactions with CrkL, which interestingly can additionally undergo CSH3-mediated homodimerization in cells.^47^ Notably, the SH2-mediated localization of CrkL at FAs is reported to be important for correct co-recruitment of ASAP1 (via the ASAP1 SH3 domain) to FAs in platelets^62^. Further studies would be needed to confirm whether PEAK3 has a role in this process also.

Finally, whilst SH3/PRM interactions do not typically require PTMs for binding, they are by no means static, and we exemplify two mechanisms by which they can be dynamically regulated: via clustering/avidity, as described above; and via accessibility, which can be modulated by the phosphorylation of an adjacent site. We identify a conserved tandem motif (CrkII^NSH3^ site / 14-3-3 site) present on all PEAKs (Fig. 5) and define a role for 14-3-3 as a potential negative regulator of PEAK3/Crk interactions at FAs (Fig. 7 and Extended Data Fig. 7a). We demonstrate that PEAK3^FL^ forms a highly stable dimer:dimer complex with 14-3-3, an interaction that prevents stable PEAK3 binding to dimeric CrkII^FL^ (Fig. 5). Our ITC competition studies with PEAK3 tandem peptides confirm that the CrkII^NSH3^ and 14-3-3 directly exhibit negative cooperativity for binding to the PEAK3 tandem site, providing a mechanistic explanation for the observed effect on PEAK3/CrkII interaction with full-length proteins (Fig.6).

PEAK3 appears to be unique from PEAK1/2 in its interaction with 14-3-3 at the tandem motif, displaying ∼ 10 fold higher affinity for 14-3-3 (PEAK3 pS69), compared to either PEAK1 (pT1165) or PEAK2 (pS826). For PEAK3, the pS69 site is sufficient for stable 14-3-3 recruitment, whereas the longer IDRs of PEAK1/2 are predicted to harbor additional 14-3-3 motifs and different mechanisms may exist. For example, the conserved tandem site 14-3-3 motif might act as a phosphodependent secondary (low-affinity) 14-3-3 site to alter PEAK1/2 activity following initial 14-3-3 recruitment at a distal ‘gatekeeker’ (high-affinity) site^63,64^.

14-3-3 binding has been reported to provide a regulatory switch in proteins involved in cytoskeletal dynamics (e.g. SSH1L, cortactin and IRSp53) by regulating downstream effector interactions and subcellular localization^65-68^. For IRSp53, 14-3-3 binding masks SH3-mediated docking with effectors at lamellipodia and filopodia, terminating IRSp53 signaling^66-68^. As CrkII and Grb2 appear to cooperate in PEAK3 signalling^14^, PEAK3-pS69 phosphorylation and 14-3-3 binding may thus provide an analogous negative feedback loop to terminate PEAK3/Crk/Grb2-mediated signaling at FAs, possibly also by altering PEAK3 subcellular localization (Fig. 7, Extended Data Fig. 7). Consistent with our findings that pS69A mutation of PEAK3 disrupts 14-3-3 binding and leads to a modest increase in association with CrkII, we therefore predict that this may also enhance PEAK3 pY24 phosphorylation and Grb2/ASAP1 association, possibly in an integrin-dependent manner. We and others have reported that PEAK3 forms complexes in cells with Grb2/PYK2/ASAP1^14,23^. In THP1 cells in the absence of growth factors, PEAK3 binding was reported to result in PYK2-mediated activation of ASAP1 and activation of PI3K/AKT signalling^23^. We propose that the S69 site on PEAK3 may therefore act as a site of negative regulation of PEAK3 activity in these signaling pathways, via phosphorylation by a basophilic serine/thronine kinase, however further studies will be required to are required to confirm the relevant kinase/s for the PEAK3 S69 site in cells.

## Conclusion

Together our work highlights distinct mechanisms of regulation and signaling outputs possible from homo/heterodimeric PEAK complexes (Extended Data Fig. 7b). We characterize the role of the conserved IDR motif and dimerisation/avidity in CrkII binding, features expected to be common to all PEAK complexes. We illuminate additional features of an IDR interaction with Grb2 present only in PEAK3/PEAK1. Lastly, we identify a role for the IDR tandem site in mediating PEAK binding to 14-3-3 proteins. For PEAK3, phosphoregulation at S69 enables formation of a distinct high affinity PEAK3:14-3-3 complex and we propose a mechanistic role in negative regulation of PEAK3/CrkII signaling. This provides a framework to better understand the contribution of intrinsically disordered regions to signaling of dimeric PEAK family pseudokinase scaffolds in healthy cells and cancer.

## Illustrations

All illustrations were generated using PyMOL, UCSF ChimeraX, PRISM, Microsoft Excel or Adobe Illustrator and InDesign.

## Reporting Summary

Further information on research design is available in the Nature Research Reporting Summary linked to this article.

## Data Availability

Coordinates and structure factors for the X-ray crystal structures have been deposited in the PDB with accession codes 8DGO (Grb2^FL^:PEAK SH2-pY peptide), 8DGP (14-3-3ε:PEAK3^tandem^-pS69), 8DGM (14-3-3ε:PEAK1^tandem^-pT1165) and 8DGN (14-3-3ε: PEAK2^tandem^-pS826).

## Acknowledgments

We thank the scientific and technical assistance of Monash Proteomics and Metabolomics, Monash University, Victoria, Australia and specifically Dr David Steer. We would like to thank the staff at the Bio21 C3 Collaborative Crystallization Centre where all initial crystallization experiments were performed and the beamline staff at the Australian Synchrotron where diffraction data were collected. **Funding:** We are grateful to the National Health and Medical Research Council (APP1144149) and Australian Research Council (DP190103672 and DP220103638) for Grant support with additional support from the Operational Infrastructure Support Program provided by the Victorian Government, Australia and from the Australian Cancer Research Foundation (to M.J.R., M.S., W.D., J.M.H., A.K., L.Y.L, O.P., I.S.L.). J.M.H was supported by an NHMRC Investigator Grant (2008096). We acknowledge Melbourne Research Scholarship support for A.K. from the University of Melbourne. The contents of this published material are solely the responsibility of the individual authors and do not reflect the views of the NHMRC or funding partners.

## Author contributions

M.J.R, M.G.S., W.D, O.P and A.K. designed, generated and biochemically characterized proteins and their complexes. M.J.R. and M.G.S. undertook crystallization experiments and M.J.R. solved crystal structures. M.J.R. and A.K. designed experiments and generated SPR and ITC data. T.C., O.P., M.J.R., B.L. conducted and analyzed SEC-MALS experiments. M.G.S., M.J.R, O.P., T.D., L.Y.L generated mass spectrometry data. J.H, X.M., R.J.D. designed, undertook, and analyzed cell-based experiments. M.J.R, M.G.S., J.H., O.P., R.J.D. and I.S.L. analyzed all the data and wrote the manuscript with input from all authors.

## Online content

### Methods

#### PEAK Multiple Sequence Alignment (MSA) and SLiM analysis

Vertebrate orthologs of human PEAK-family proteins (PEAK3, PEAK1 and PEAK2/PRAG1) were compiled from the NBCI Gene database^1^ (calculated by NCBI’s Eukaryotic Genome Annotation pipeline) and manually curated to remove sequences annotated by NCBI as low quality. This yielded orthologs for PEAK3 (149 vertebrate orthologs), PEAK1 (301 vertebrate orthologs) and PEAK2 (206 vertebrate orthologs). Multiple Sequence Alignment for each set of PEAK orthologs and for the full family of all PEAK orthologs was performed using PROMALS3D^2^ and visualized by web logo generation using WebLogo3^3^. Short Linear Motif (SLiM) analysis was performed using the Eukaryotic Linear Motif (ELM) server^4^ and Scansite 4.0^5^ and manual annotation.

#### Synthetic peptides and commercial recombinant protein

All synthetic peptides were purchased from Mimotopes (95% purity).

##### Proline rich motif peptides

13-mer peptides encompassing proline-rich motifs from PEAK proteins were synthesized: PEAK3^54-66^ (Ac-PLPPPLPKKILTR-NH2); PEAK1^1117-1129^ (Ac-MIPPKQPRQPKGA-NH2); PEAK1^1150-1162^ (Ac-PTPPPLPKKMIIR-NH2); PEAK2^709-721^ (Ac-FSPPPPPPKSRHL-NH2); and PEAK2^809-821^ (Ac-QPPPLPQKKIVSR-NH2).

##### SH2 peptides

A 7-mer sequence containing the putative Grb2/CrkII SH2 binding motif is common to both human PEAK3 (residues 23-29) and human PEAK1 (residues 1106-1112). Two synthetic 7mer peptides corresponding to this sequence were synthesized, either phosphorylated or non-phosphorylated at the tyrosine residue (PEAK SH2-pY peptide, Ac-T(pY)SNLGQ-NH2; and Ac-TYSNLGQ-NH2; both N-terminal acetylated, C-terminally amidated)^6^.

##### Tandem peptides

Tandem peptides (phosphorylated and non-phosphorylated at the bolded/underlined residue) were also synthesized: PEAK3^tandem^ and PEAK3^tandem^-pS69 (residues 54-74; Ac-PLPPPLPKKILTRTQSLPTRR-NH2); PEAK1^tandem^ and PEAK1^tandem^-pT1165 (residues 1152-1171; Ac-PPPLPKKMIIRANTEPISKD-NH2); and PEAK2^tandem^ and PEAK2^tandem^-pS826 (residues 812-831; Ac-PPPLPQKKIVSRAASSPDGF-NH2).

#### Protein production of His-tagged adapter/scaffold proteins

Human gene sequences of adapter/scaffold proteins (full length or truncated versions) were codon optimized for *E. coli* expression and cloned into a pCOLD-IV vector (Takara) modified to encode an N-terminal 8-His tag and tobacco etch virus (TEV) protease cleavage site for tag removal. Several constructs were generated for this study. For adapter proteins this included: full length Grb2 (Grb2^FL^; UniProt P62993-1, residues 1-217); full length CrkII (CrkII^FL^; UniProt P46108-1, residues 1-330), and truncated forms of CrkII, including a SH2-NSH3 domain construct lacking the CSH3 domain (CrkII^ΔCSH3^ residues 1-228) and constructs of the individual CrkII SH2 domain (CrkII^SH2^; UniProt P46108-1, residues 6-120) and CrkII NSH3 domain (CrkII^NSH3^; UniProt P46108-1, residues 134-191). For 14-3-3 isoforms this included: 14-3-3ε (UniProt P62258-1; residues 1-255), 14-3-3η (UniProt Q04917-1; residues 1-246), 14-3-3γ (UniProt P61981-1; residues 1-247) and 14-3-3σ (UniProt P63104-1; residues 1-245). All constructs were verified using Sanger Sequencing (Micromon). Expressions were carried out overnight at 16°C for 16 to 20 hours in *E. coli* C41 (DE3) and proteins were purified using nickel affinity chromatography and size exclusion chromatography (SEC). When required, the His-tagged protein was cleaved following SEC with TEV protease. Cleaved samples were further purified by ion exchange chromatography (IEX) using Mono Q™ 5/50 GL (CrkII^FL^, 14-3-3 and Grb2^FL^) columns (Cytiva) (Extended Data Table 1). All protein samples were concentrated, flash frozen and stored at -80°C.

#### Protein production of PEAK proteins

Human PEAK1 (PEAK1^IDR1^; UniProt Q9H792, residues 1082-1746), PEAK2 (PEAK2^IDR1^; UniProt Q86YV5, residues 802-1406) and PEAK3 (PEAK3^FL^; UniProt Q6ZS72, residues 1-473) were codon optimized for insect cell expression and cloned into a modified pFastBac*™* Dual vector containing a N-terminal TEV cleavable His-tag under the polyhedrin promoter^7^. Expression was carried out in *Spodoptera frugiperda* (*Sf*21) cells and purification was performed as previously described^7^ (Extended Data Table 2). Briefly, cell lysates were clarified by centrifugation at 45,500 x g for 1 hour at 4 °C and mixed with 0.5% (v/v) of His-tag purification resin (Roche) for purification by nickel affinity chromatography, PEAK proteins were further purified by SEC (HiLoad 16/600 Superdex 200 pg column). For PEAK3^FL^, the resulting sample contained a mixture of PEAK3 protein in complex with two proteins identified by mass spectrometry as endogenous insect cell-derived 14-3-3ζ and 14-3-3ε (Extended Data Fig. 4a). PEAK1^IDR1^ and the PEAK3^FL^:14-3-3ε,ζ complex were further purified by IEX (Mono Q™ 5/50 GL column). PEAK1^IDR1^, PEAK2^IDR1^ and the PEAK3^FL^: 14-3-3ε,ζ complex were concentrated to ∼ 5 mg ml^−1^ and flash frozen in liquid nitrogen for subsequent studies.

#### Recombinant kinase domains for *in vitro* phosphorylation reactions

Constructs for expression of recombinant Src kinase domain (UniProt P12931, residues 254-536; Addgene plasmid #79700), Abl kinase domain (UniProt P00519-1, residues 229-512; Addgene plasmid #78173) and untagged YopH phosphatase (UniProt P08538-1, residues 164-468; Addgene plasmid #79749) from the Human Kinase Domain Constructs Kit for Automated Bacterial Expression (Addgene plasmid kit #1000000094) were a gift from John Chodera (Memorial Sloan Kettering Cancer Center)^8^. Co-expressions of Src/YopH or Abl/YopH were carried out overnight at 16°C for 16 to 20 hours in *E. coli* C41 (DE3) or *E. coli* Rosetta (DE3), respectively and purified as described using nickel affinity chromatography and SEC, without cleavage of the His-tag. Purified Abl kinase domain was stored at 2 mg ml^−1^ in 20 mM Tris pH 7.5, 150 mM NaCl, 5% glycerol, 0.5 mM TCEP. The Src kinase domain was further purified by IEX (Mono Q™ 5/50 GL column) and stored at 12 mg ml^−1^ in 20 mM Tris pH 8.0, 165 mM NaCl, 1 mM TCEP for subsequent studies (Extended Data Table 3).

#### Complex studies using analytical SEC and SEC-MALS

##### SEC analysis of protein complexes

SEC analysis of PEAK3^FL^, PEAK2^IDR1^, PEAK1^IDR1^ and CrkII^FL^ dimer/monomer (Fig. 3-4 and Extended Data Fig. 3) were performed on a Superdex™ 200 Increase 10/300 GL (GE Healthcare) equilibrated in 20 mM HEPES pH 7.5-8.5, 150 mM NaCl, 1 mM TCEP. For individual analysis, PEAK and CrkII proteins were prepared at 20-50 μM concentration in a total volume of 110 μl. For complex interaction studies, in general PEAK and CrkII proteins were mixed at a 1:1-2 molar ratio (20-50 μM final concentration) in a total reaction volume of 110 μl, and the complex pre-equilibrated at room temperature for 25 min and filtered prior to SEC analysis. Elution peak fractions were analyzed by SDS-PAGE and peak elution fractions were pooled and concentrated and flash frozen in liquid nitrogen for subsequent studies.

##### SEC-MALS analysis of protein complexes

SEC multi-angle light scattering (SEC-MALS) experiments were performed using a Superdex 200 Increase 10/300 GL column (Cytiva) coupled with a DAWN HELEOS II light scattering detector and an Optilab TrEX refractive index detector (Wyatt Technology). The system was equilibrated in 20 mM HEPES pH 7.5, 150 mM NaCl, 1 mM TCEP and calibrated using bovine serum albumin (2 mg ml^−1^; Sigma) before analysis of experimental samples. For each experiment, 100 µl of purified pre-equilibrated protein complex (120 μg of PEAK3^FL^: 14-3-3ε,ζ and 170 μg of PEAK2^IDR1^:CrkII^FL^ dimer) was injected onto the column and eluted at a flow rate of 0.5 ml min^−1^. Experimental data were collected and processed using ASTRA software (Wyatt Technology, v.7.3.19).

#### Mass spectrometry (MS)

##### Tryptic MS phosphosite analysis of recombinant PEAK proteins

Recombinant purified PEAK1^IDR1^, PEAK2^IDR1^ and PEAK3^FL^ were reduced with DTT (Sigma), carbamidomethylated with iodoacetamide (Sigma), and digested with trypsin (Promega). Prior to MS, the peptides were further purified and enriched using OMIX C18 Mini-Bed tips (Agilent Technologies). Using a Dionex UltiMate 3000 RSLCnano system equipped with a Dionex UltiMate 3000 RS autosampler, an Acclaim PepMap RSLC analytical column (75 μm × 50 cm, nanoViper, C18, 2 μm, 100 Å; Thermo Fisher Scientific), and an Acclaim PepMap 100 trap column (100 μm × 2 cm, nanoViper, C18, 5 μm, 100 Å; Thermo Fisher Scientific), the tryptic peptides were separated by increasing concentrations of 0.1% (v/v) formic acid (prepared in 80% (v/v) acetonitrile) at a flow rate of 250 nl min^−1^ for 50 min and analyzed with a QExactive HF mass spectrometer (Thermo Fisher Scientific).

To obtain peptide and phosphopeptide sequence information, the raw files were searched with Byonic (Protein Metrics, v. 3.1.0) against a human database obtained from UniProt (UP000005640) with the following parameters: carbamidomethylation at cysteine residues was specified as a fixed modification; oxidation at methionine residues and phosphorylation at serine, threonine, or tyrosine residues were set as variable modifications; cleavage site specificity was set at C-terminal residues K and R with semi-specific cleavage specificity with up to two missed cleavages permitted; and precursor and fragment ion mass tolerances were set to 20 ppm. Only peptides identified within a false discovery rate of 1% based on a decoy database were reported.

##### In vitro ABL/SRC phosphorylation of PEAKs, CrkII and PTM analysis using tandem MS/MS

Phosphorylation reactions – Purified recombinant PEAK3^FL^ and PEAK1^IDR1^, CrkII^FL^ were phosphorylated *in vitro* using recombinant Abl and Src. Substrate proteins were incubated at room temperature for 3 hours with recombinant Abl or Src kinase domains (5:1 molar ratio of substrate to kinase, 100 µl reaction volume) in 40 mM Tris pH 8.0, 150 mM NaCl, 10 mM MgCl2, 1 mM DTT and 5 mM ATP. Reactions were stopped with 25 mM EDTA. Control samples (no Abl or Src added to reaction) were performed analogously and included in MS analysis. MS analysis was performed as for recombinant PEAK proteins. Peptide and phosphopeptide sequences were obtained in Byonic using the same parameters as the tryptic MS phosphosite analysis.

#### Protein crystallization, data collection and processing

##### Grb2^FL^:PEAK SH2-pY peptide complex

Crystallization of the Grb2^FL^:PEAK SH2-pY peptide complex was accomplished based on the previously described method for Grb2^FL^ alone^9^, using the hanging-drop vapor diffusion method at 20°C, and initial crystals optimized by varying pH, streak seeding and adjusting the drop ratio. The protein complex was prepared by concentrating Grb2^FL^ to 16 mg ml^−1^ with 1 mM peptide (diluted from 10 mM stock in water). Crystals used for data collection were obtained in 2.6 M sodium acetate pH 8.0, by mixing the protein/peptide complex with the well solution in a 2:1 ratio. Crystals were cryo-protected in 2.7 M sodium acetate pH 8.0, 10 % (v/v) glycerol, supplemented with 1 mM peptide and flash-frozen in liquid nitrogen. Diffraction data were collected on beamline MX2 at the Australian Synchrotron^10^ using a wavelength of 0.9537 Å. Data were integrated by using XDS (v. 20161205)^11^, scaled by using XSCALE (v. 20161205), and merged using AIMLESS (v. 0.5.21)^12,13^. The crystals belonged to space group *P* 4_3_ with unit cell parameters a = b = 89.0, c = 94.5 Å and α = β = γ = 90°, and contained two copies of Grb2^FL^ per asymmetric unit as a closely packed homodimer. The structure was solved by molecular replacement using PHASER (v. 2.8.3)^14^ with Grb2^FL^ monomer coordinates derived from the existing Grb2^FL^ structure (PDB entry 1GRI) as the initial search model. Clear unbiased density was observed for the phosphopeptide in the SH2 phosphotyrosine binding site for one copy of Grb2^FL^ in the dimer, whilst no additional density was observed in the SH2 site in the second copy, which appeared to be occluded due to crystal packing. Iterative model building and refinement were performed in COOT (v. 0.9.8.1)^15^ and PHENIX (v. 1.20.1-4487)^16^, respectively, including a simulated annealing step in the first round of refinement to minimize model bias and manual building of the phosphopeptide. The structure was refined to Rwork and Rfree values of 16.6% and 20.9%, respectively, with all of the residues in Ramachandran allowed regions as validated by MOLPROBITY^17^. The overall structure aligns closely with the earlier published structure for Grb2^FL^ alone (PDB entry 1GRI, 3.1 Å resolution), with the addition of the PEAK1/3 pY-site phosphopeptide and some additional Grb2^FL^ residues able to be modelled due to the higher resolution (2.3 Å resolution).

##### 14-3-3ε:PEAK3^tandem^-pS69 complex

The 14-3-3ε:PEAK3^tandem^-pS69 peptide complex was prepared by incubating 14-3-3ε (10 mg ml^−1^ final concentration) with the PEAK tandem peptide (0.5 mM final concentration, diluted from 10 mM stock in water), representing a 1.5-fold molar excess of peptide. The complex was subjected to sparse matrix screening in a 96-well plate using the sitting-drop vapor diffusion method at 20°C and a 1:1 drop ratio of protein and well solution. Diffracting crystals were obtained in 2 M ammonium sulfate, 0.1 M sodium bicine pH 9.35, and 5% (v/v) 2-methyl-2,4,-pentandiol (MPD). Crystals were cryo-protected in 2.2 M ammonium sulfate, 0.02 M sodium bicine pH 9.35, 5% (v/v) MPD, and 20% (v/v) ethylene glycol supplemented with 100 µM peptide and flash-frozen in liquid nitrogen. Data collection and processing was performed as for the Grb2^FL^:PEAK SH2-pY peptide complex. The crystals belonged to space group P 4_3_ with unit cell parameters a = b = 155.42, c = 58.0 Å and α = β = γ = 90°, and contained four copies of 14-3-3ε monomer per asymmetric unit, arranged as two symmetrical homodimers as anticipated. Molecular replacement, model building and refinement were performed as for the Grb2^FL^:PEAK SH2-pY peptide complex except that the human 14-3-3ε monomer structure (PDB entry 2BR9) was used as the initial search model. Clear unbiased density was observed for the phosphopeptide in the binding site of each 14-3-3ε monomer, although chain A showed the clearest resolved density following building and refinement. The structure was refined to Rwork and Rfree values of 18.7% and 23.3%, respectively, with 99.9% of the residues in Ramachandran allowed regions as validated by MOLPROBITY^17^.

##### 14-3-3ε:PEAK1^tandem^-pT1165 complex

The structure of the 14-3-3ε:PEAK1^tandem^-pT1165 complex was obtained by firstly crystallizing 14-3-3ε alone and then soaking the crystals with the PEAK tandem phosphopeptide. Successful conditions identified for the 14-3-3ε:PEAK3^tandem^-pS69 peptide complex were used as the basis for 14-3-3ε crystallization trials using the hanging-drop vapor diffusion method at 20°C. Crystals of 14-3-3ε (10 mg ml^−1^) were obtained in 1.8 M ammonium sulfate, 0.1 M HEPES pH 7.5, 5% (v/v) MPD, then transferred to well solution supplemented with 0.5 mM phosphopeptide to soak overnight. Crystals were cryo-protected in 2.2 M ammonium sulfate, 0.1 M HEPES pH 7.5, 5% (v/v) MPD, 20% (v/v) ethylene glycol supplemented with 0.5 mM phosphopeptide and flash-frozen in liquid nitrogen. Data collection, molecular replacement, and model building and refinement were performed as for the 14-3-3ε:PEAK3^tandem^-pS69 complex. The crystals belonged to space group *P* 6_2_ 2 2 with unit cell parameters a = b = 91.4, c = 139.9 Å and α = β = 90°, γ = 120° and contained one 14-3-3ε monomer per asymmetric unit. This particular crystal form represents a close crystal packing of 14-3-3ε as a monomer, in which residues 1-32 that would typically mediate the 14-3-3ε dimer interface are not resolved, as the crystal packing would require them to be at least partially unfolded. The remainder of the 14-3-3ε protein core aligns closely with the structure of dimeric 14-3-3ε (as for the 14-3-3ε*:PEAK3*^*tandem*^*-pS69 complex*). Notably, the peptide binding groove is accessible and amenable to soaking and clear unbiased density for the PEAK phosphopeptide was observed. The final structure was refined to Rwork and Rfree values of 20.7% and 26.6%, respectively, with 99.5% of the residues in Ramachandran allowed regions as validated by MOLPROBITY^17^.

##### 14-3-3ε:PEAK2^tandem^-pS826 complex

Crystallization, soaking, data collection and processing, and model building and refinement were performed identically as for the 14-3-3ε:PEAK1^tandem^-pT1165 complex, but using the PEAK2^tandem^-pS826 phosphopeptide. The crystals belonged to space group *P* 6_2_ 2 2 with unit cell parameters a = b = 92.1, c = 137.4 Å and α = β = 90°, γ = 120°, and contained one 14-3-3ε monomer per asymmetric unit, in the same close packing of the 14-3-3ε monomer. The final structure was refined to Rwork and Rfree values of 24.2% and 29.5%, respectively, with all the residues in Ramachandran allowed regions as validated by MOLPROBITY^17^.

#### AlphaFold2 structure prediction of PEAK3^FL^ homodimer

The ColabFold Google Colab notebook called AlphaFold2_complexes^18,19^ was used to predict the structure of dimeric human PEAK3^FL^. The predicted model with the highest lDDT score is shown.

#### AlphaFold2 / HADDOCK analysis of SH2-pY motif

The PEAK3^23-27^-pY^24^ peptide (T(pY)SNL) from the crystal structure of the Grb2^FL^:PEAK SH2-pY peptide complex was spliced with the predicted structure of human PEAK3^28-38^ (GQIRAHLLPSK) from the AlphaFold Protein Structure Database (EMBL-EBI)^20^. The spliced peptide was then docked into the Grb2^FL^ crystal structure using the HADDOCK server (v. 2.4)^21^ and the highest ranked model was analyzed in ChimeraX (v. 1.4)^22^ (Extended Data Fig. 2c, Fig. 2e).

#### Surface Plasmon Resonance (SPR) binding assays

All SPR binding studies were performed using a Biacore S200 Instrument (Cytiva).

##### SPR binding studies to CrkII^NSH3^

Purified CrkII^NSH3^ was diluted to 5 µg ml^−1^ in 10 mM sodium acetate pH 4.0 and amine coupled at 25°C to a Series S CM5 sensor chip, in HBS-N running buffer (20 mM HEPES pH 7.4, 150 mM NaCl) to a final immobilization level of 1100-1400 response units (RU), followed by surface deactivation using 1 M ethanolamine. A blank activation/deactivation was used for the reference surface. Peptide binding studies were performed at 20°C in HBS-TP running buffer (20 mM HEPES pH 7.4, 150 mM NaCl, 1 mM TCEP, 0.005% (v/v) Tween-P20). Peptides were first prepared as 10 mM stocks in water: PEAK PRM peptides: PEAK3^54-66^, PEAK1^1150-1162^, PEAK2^709-721^, PEAK2^809-821^; or PEAK tandem motif peptides: PEAK3^tandem, 54-74^, PEAK3^tandem, 54-74^-pS69, PEAK1^tandem, 1152-1171^, PEAK1^tandem, 1152-1171^-pT1165, PEAK2^tandem, 812-831^, and PEAK2^tandem, 812-831^-pS826, phosphorylated or non-phosphorylated in the 14-3-3 motif. Peptide stocks were further diluted to in running buffer to 10 µM or 20 µM and prepared as a 11-point concentration series (2-fold serial dilution, 100 µM - 100 nM for PEAK PRM peptides, or 20 µM - 20 nM for PEAK tandem motif peptides). Samples were injected in a multi-cycle run (flow rate 30 µl min^−1^, contact time of 60 seconds, dissociation 120 seconds) without regeneration. Sensorgrams were double referenced, and steady state binding data fitted using a 1:1 binding model using Biacore S200 Evaluation Software (Cytiva, v. 1.1). Representative sensorgrams and fitted dissociation constant (*K*_D_) values, depicted as mean ± S.E.M. (N > 3 independent experiments) or mean ± S.D. (N = 2), are shown in Source Data Fig. 3.

##### SPR binding studies to 14-3-3 isoforms

Immobilization of 14-3-3 isoforms (14-3-3γ, 14-3-3ε, 14-3-3σ, and 14-3-3η; diluted to 10 µg ml^−1^ in 10 mM sodium acetate pH 4.0) was performed as described for CrkII^NSH3^ at 20°C to a final immobilization level 1800 – 2900 RU. Binding studies were run using PEAK tandem peptides (11-point concentration series, 2-fold serial dilution; 10 µM - 10 nM for PEAK3^tandem^-pS69, 20 µM – 20 nM for all other peptides). SPR binding experiments and steady state data analysis were performed as described for CrkII^NSH3^. Representative sensorgrams and fitted dissociation constant (*K*_D_) values, depicted as mean ± S.E.M. (N > 3 independent experiments), are shown in Source Data SPR Fig.4.

#### Isothermal Titration Calorimetry (ITC) binding experiments and cooperativity analysis

ITC binding experiments were conducted using a MicroCal iTC200 instrument (Malvern Instruments). Titrations were conducted at 25 °C in reverse mode (peptide in cell, protein in syringe) and consisted of 19 injections of 2 μL peptide solution at a rate of 2 sec μL^−1^ at 90 sec time intervals, whilst stirring at 1000 rpm. An initial injection of 0.4 μL protein was made and discarded during data analysis. Proteins (14-3-3ɣ or CrkII^NSH3^) were first dialyzed overnight at 4°C into ITC buffer (20 mM HEPES pH 7.4, 150 mM NaCl, 1 mM TCEP), filtered and degassed under vacuum. Peptides (PEAK3^tandem^ or PEAK3^tandem^pS69) were diluted into filtered, degassed ITC buffer from 10 mM stock solutions prepared in water. For the binary experiment of CrkII^NSH3^ binding to PEAK3^tandem^ peptide, CrkII^NSH3^ (140 μM, in the syringe) was titrated into PEAK3^tandem^ peptide (20 μM, in the cell). For binding studies to examine cooperativity, sequential ITC titrations were performed according to the following general procedure: In the first titration (binary), 14-3-3ɣ (200 μM, in the syringe) or a buffer control were titrated into PEAK3^tandem^ or PEAK3^tandem^pS69 peptide (20 μM, in the cell). Following the first titration, excess solution was removed from the cell, and the syringe was washed and dried. In the second titration (ternary), CrkII^NSH3^ (400 μM, in the syringe) was titrated into the peptide complex remaining in the cell (16.8 µM). The concentration of saturated peptide complex in the cell (C) after the first titration (16.8 μM), was calculated using the equation: C = (C_0_ · V_cell_)/(V_cell_+V_inj_) where C_0_ is the initial peptide concentration (20 µM), V_cell_ is the volume of the cell (200.1 µl) and V_inj_ is the volume of titrant injected during the first titration (38.4 µl). All data were fitted to a single binding site model to obtain the stoichiometry (N), the dissociation constant (*K*_D_) and the enthalpy of binding (ΔH), using Microcal Origin (OriginLab, v. 7.0). The reported values are the mean ± S.E.M. (N > 3) from independent measurements (Extended Data Fig. 6).

#### Cellular studies of PEAK interactions (HEK293 cells)

##### Antibodies and tissue culture

The following antibodies were obtained commercially: anti-HA (Cell Signaling Technology, catalog no. 3724), anti-Flag (Sigma-Aldrich, catalog no. F1804), anti-14–3-3 (Santa Cruz Biotechnology, catalog no. sc-1657) and anti-CrkII (Santa Cruz Biotechnology, catalog no. sc-289). HEK293 cells were maintained as previously described^6^.

##### Plasmids

Codon-optimized cDNAs encoding N-terminal HA-tagged or Flag-tagged WT PEAK3^FL^ and HA-tagged PEAK3^FL^ mutants were synthesized as previously described^6^. Plasmid transfections in HEK293 cells were performed using Lipofectamine 3000 (Life Technologies, catalog no. L3000015) according to the manufacturer’s instructions.

##### Cell lysis and immunoprecipitation

Control HEK293 cells or HEK293 cells singly expressing HA-tagged PEAK3^FL^ WT or HA-tagged PEAK3^FL^-S69A mutant, or co-expressing Flag-tagged PEAK3^FL^ WT with various HA-tagged PEAK3^FL^ dimerisation mutants (PEAK3^FL^-S69A, -L146E, -A436E, or -C453E) were harvested after 48h post transfection. Cell lysates for co-immunoprecipitation were prepared using standard lysis buffers^23^. Cell lysates were incubated with anti-HA affinity-agarose beads (Sigma-Aldrich, catalog no. E6679) at 4°C overnight on a shaking platform. After extensive washing with ice-cold lysis buffer, the immune complexes were eluted with SDS–polyacrylamide gel electrophoresis loading buffer and subjected to Western blotting analysis. Uncropped gels can be found in Source data.

---

**Extended Data Table 1.** Purification buffers for adaptor/scaffold proteins.

| Buffers | 14-3-3 isoforms | CrkII<sup>FL</sup><br>CrkII<sup>ΔCSH3</sup><br>CrkII<sup>NSH3</sup><br>CrkII<sup>SH2</sup> | Grb2<sup>FL</sup><br>Grb2<sup>SH2</sup> |
| --- | --- | --- | --- |
| Lysis | 20 mM Tris pH 7.5<br>500 mM NaCl<br>5 mM DTT<br>5 mM imidazole<br>10% (v/v) glycerol<br>0.1% (w/v) Thesit<br>2 mM EDTA | 20 mM Tris pH 7.5<br>500 mM NaCl<br>5 mM DTT<br>5 mM imidazole<br>10% (v/v) glycerol<br>0.1% (w/v) Thesit | 20 mM Tris pH 7.5<br>500 mM NaCl<br>5 mM DTT<br>5 mM imidazole<br>10% (v/v) glycerol<br>0.1% (w/v) Thesit |
| Ni-Affinity Wash | 20 mM Tris pH 7.5<br>500 mM NaCl<br>5 mM DTT<br>5 mM imidazole<br>10% (v/v) glycerol<br>2 mM EDTA | 20 mM Tris pH 7.5<br>500 mM NaCl<br>5 mM DTT<br>5 mM imidazole<br>10% (v/v) glycerol | 20 mM Tris pH 7.5<br>500 mM NaCl<br>5 mM DTT<br>5 mM imidazole<br>10% (v/v) glycerol |
| Ni-Affinity Elution | Ni-Affinity Wash Buffer<br>250 mM imidazole | Ni-Affinity Wash Buffer<br>250 mM imidazole | Ni-Affinity Wash Buffer<br>250 mM imidazole |
| SEC | 20 mM HEPES pH 7.5<br>150 mM NaCl<br>0.5 mM TCEP | 20 mM Tris pH 7.5<br>200 mM NaCl<br>0.5 mM TCEP<br>5% (v/v) glycerol | 20 mM Tris pH 7.5<br>200 mM NaCl<br>0.5 mM TCEP<br>5% (v/v) glycerol |
| IEX Buffer A | 20 mM HEPES pH 7.5<br>0.5 mM TCEP | 20 mM HEPES pH 7.5<br>0.5 mM TCEP | 20 mM HEPES pH 7.5<br>0.5 mM TCEP<br>5% (v/v) glycerol |
| IEX Buffer B | 20 mM HEPES pH 7.5<br>1 M NaCl<br>0.5 mM TCEP | 20 mM HEPES pH 7.5<br>1 M NaCl<br>0.5 mM TCEP | 20 mM HEPES pH 7.5<br>1 M NaCl<br>0.5 mM TCEP<br>5% (v/v) glycerol |

**Extended Data Table 2.**
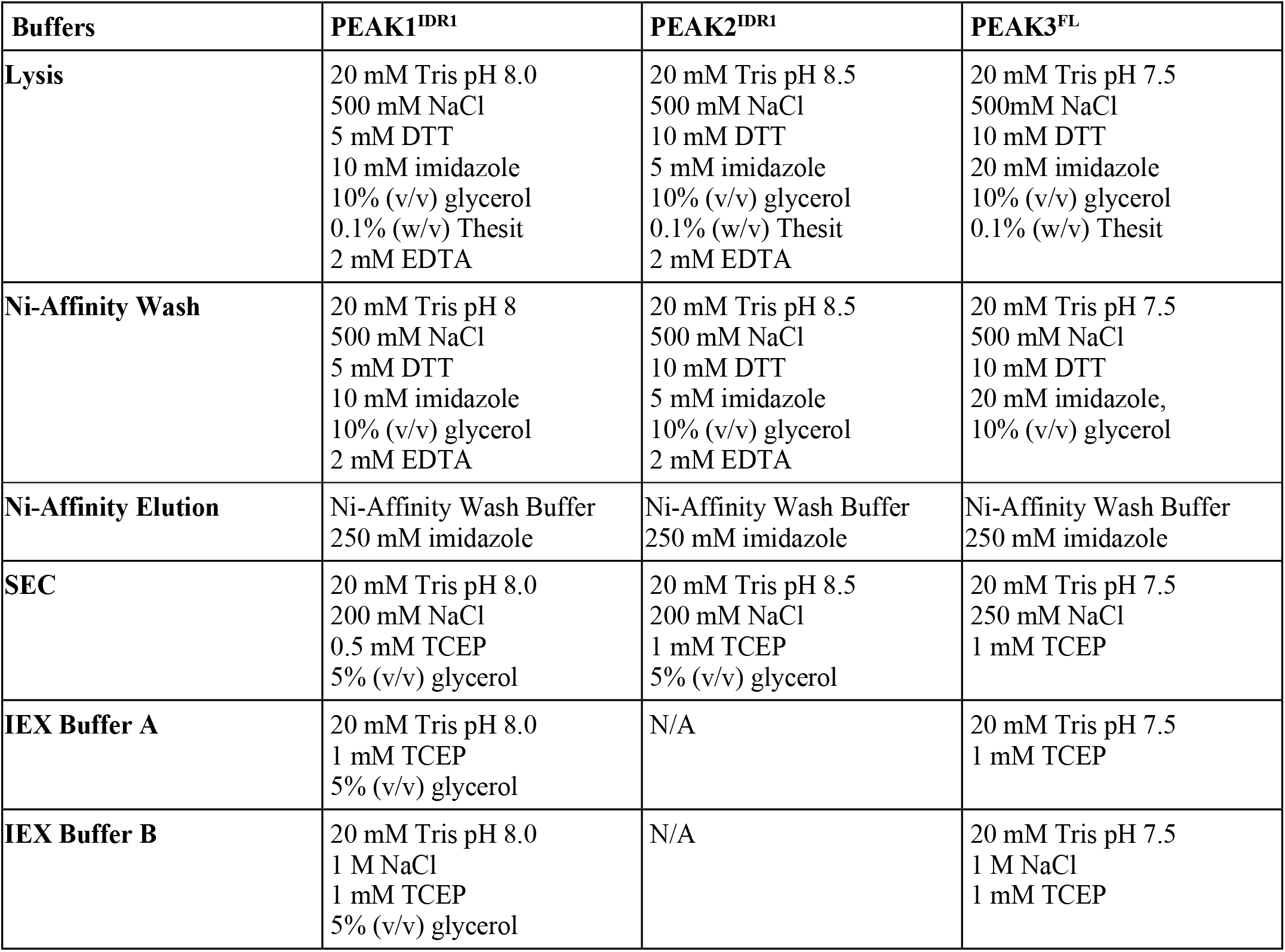
Purification buffers for PEAK proteins.

**Extended Data Table 3.** Purification buffers for kinase domains for *in vitro* phosphorylation reactions.

| Buffers | Src | Abl |
| --- | --- | --- |
| **Lysis** | 50 mM Tris pH 8.0<br>500 mM NaCl<br>1 mM DTT<br>10 mM imidazole<br>5% (v/v) glycerol<br>0.1% (w/v) Thesit | 20 mM Tris pH 7.5<br>500 mM NaCl<br>5 mM DTT<br>5 mM imidazole<br>10% (v/v) glycerol<br>0.1% (w/v) Thesit |
| **Ni-Affinity Wash** | 50 mM Tris pH 7.5<br>500 mM NaCl<br>1 mM DTT<br>25 mM imidazole<br>5% (v/v) glycerol | 20 mM Tris pH 7.5<br>500 mM NaCl<br>5 mM DTT<br>5 mM imidazole<br>10% (v/v) glycerol |
| **Ni-Affinity Elution** | Ni-Affinity Wash Buffer<br>250 mM imidazole | Ni-Affinity Wash Buffer<br>250 mM imidazole |
| **SEC** | 20 mM Tris pH 8.0<br>150 mM NaCl<br>1 mM DTT | 20 mM Tris pH 7.5<br>150 mM NaCl<br>5% (v/v) glycerol<br>0.5 mM TCEP |
| **IEX Buffer A** | 20 mM Tris pH 8.0<br>1 mM TCEP | N/A |
| **IEX Buffer B** | 20 mM Tris pH 8.0<br>1 M NaCl<br>1 mM TCEP | N/A |

## Extended Data Figure Legends

**Extended Data Fig. 1.**
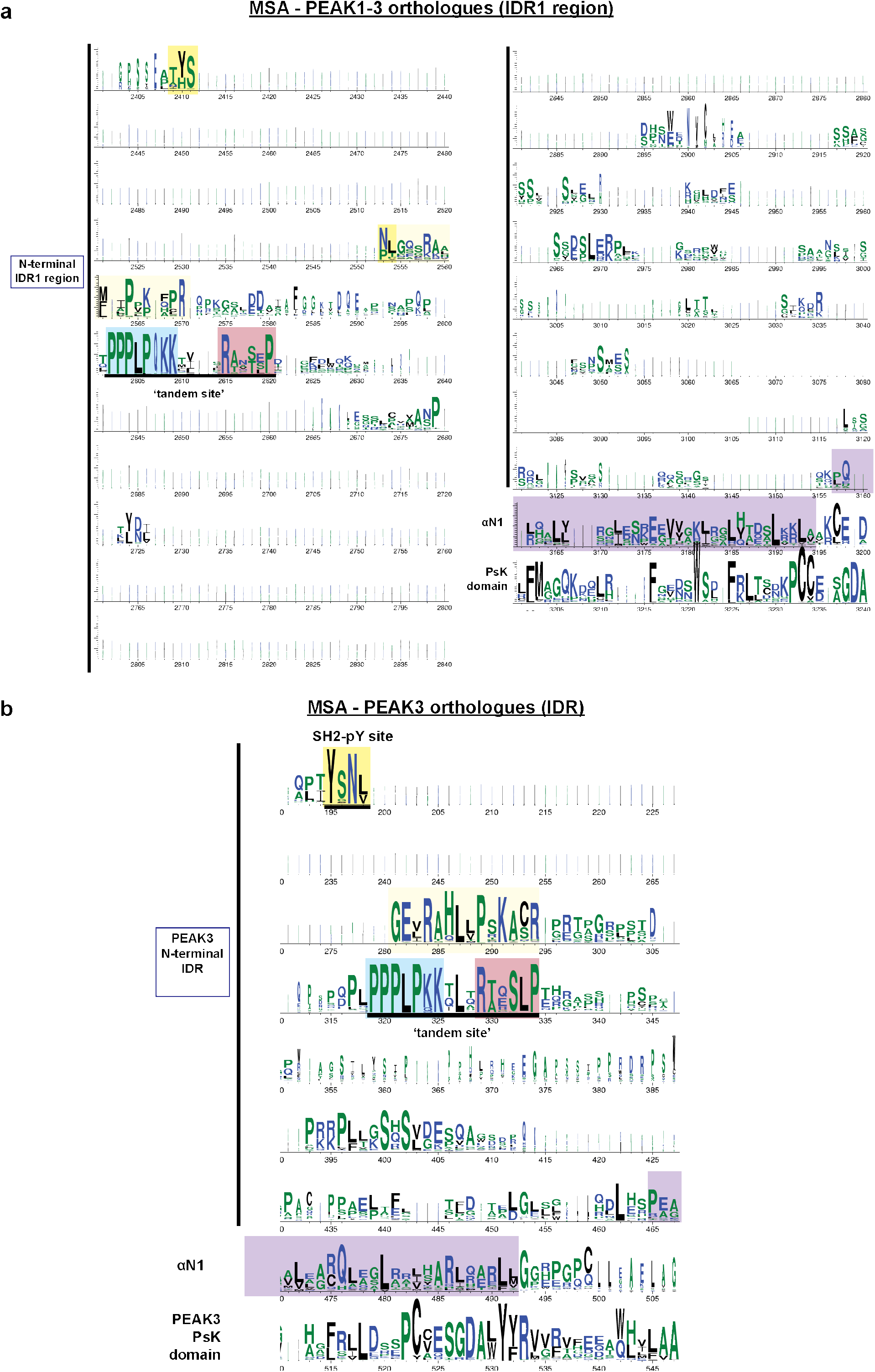
**a**, Full MSA/Web Logo of conservation across PEAK3/PEAK1/PEAK2 vertebrate orthologs (N-IDR1 region). **b**, Full MSA/Web Logo of PEAK3 orthologs (N-terminal IDR1 region). Regions of interest for this study are highlighted, in particular the conserved SH2 pY binding site present in PEAK3/PEAK1 (TYSNL motif) and the ‘tandem site’ (encompassing proline rich motif known to bind CrkII^NSH3^ and putative 14-3-3 binding motif).

**Extended Data Fig. 2.**
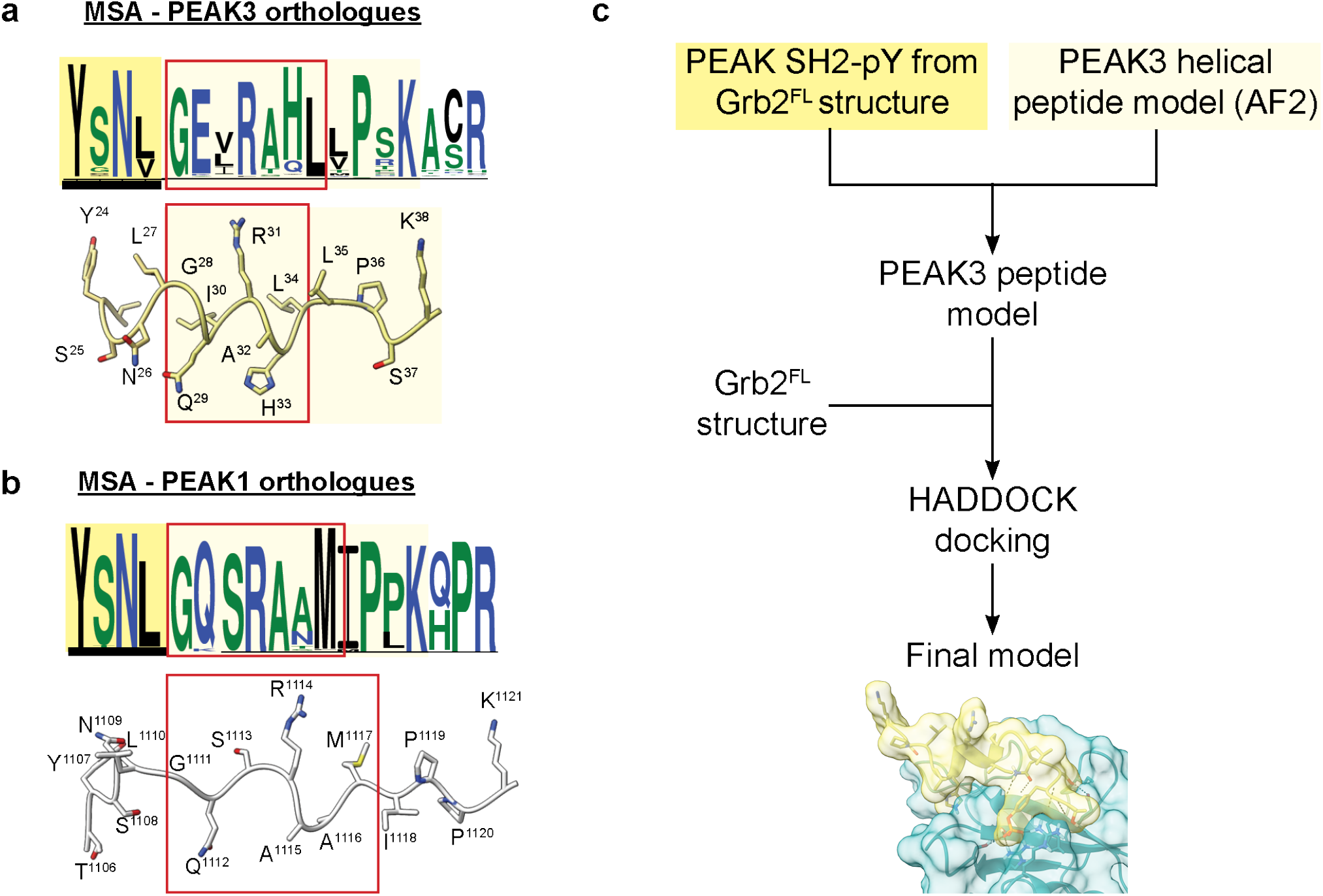
**a-b**, AlphaFold2 (AF2) structures of (a) PEAK1^1106-1121^ and (b) PEAK3^23-38^ regions alongside Full MSA/Web Logo of orthologs for the corresponding sequence, showing a section of high sequence conservation (pale yellow box) adjacent to the TYSNL SH2 site (yellow box, underlined) including a region predicted in AF2 to adopt helical secondary structure (red box). **c**, Workflow to generate HADDOCK docking model of extended PEAK3^23-38^-pY^24^ peptide (TpYSNL motif and predicted helical region) bound to Grb2^FL^.

**Extended Data Fig. 3.**
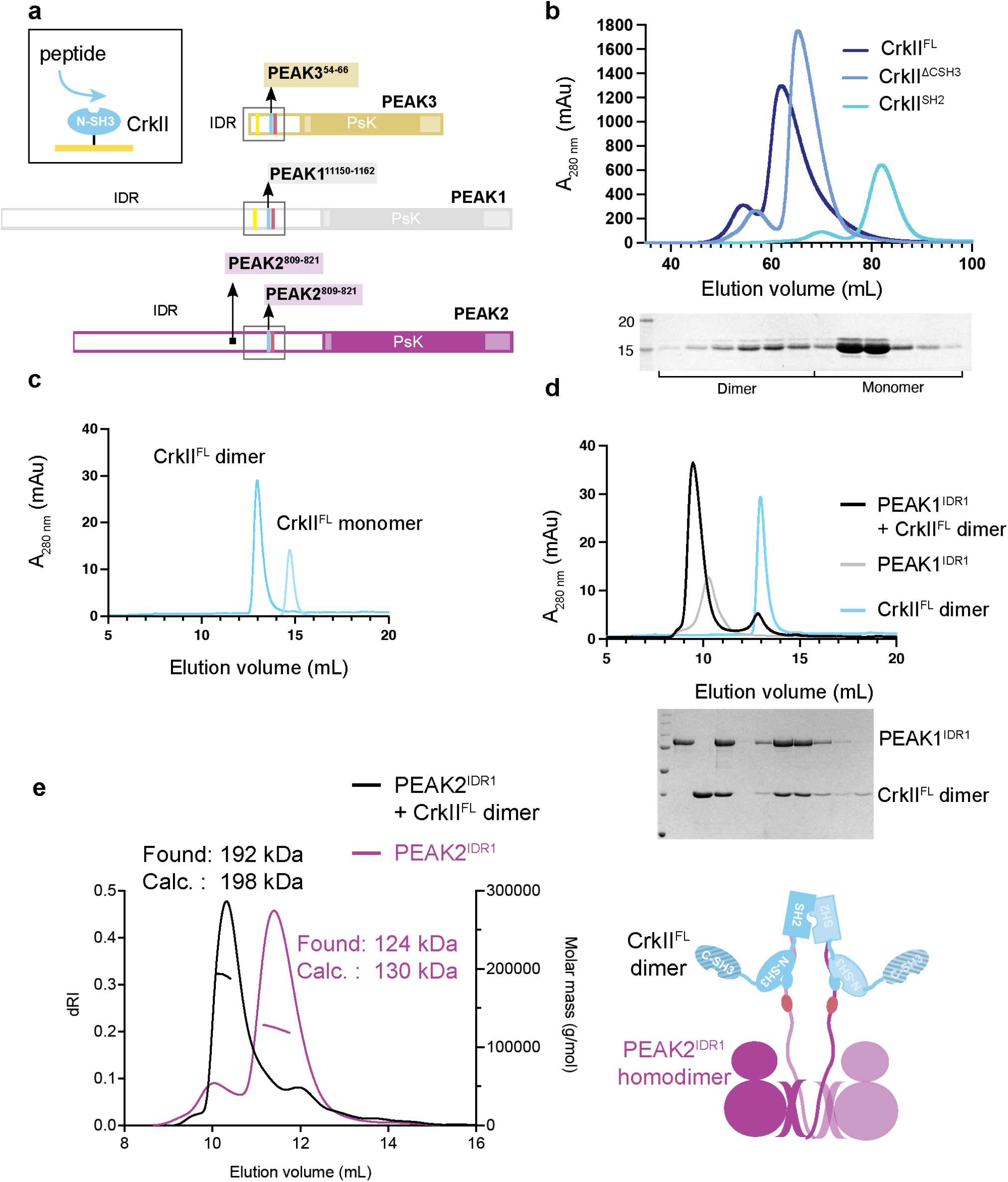
**a**, Schematic highlighting synthetic peptides generated for PEAK1, PEAK2 or PEAK3 encompassing Proline Rich Motifs (PRMs) predicted to bind CrkII^NSH3^ domain. **b**, Size exclusion chromatography (SEC) profile (S75 16/600) of CrkII^FL^ and truncated CrkII^ΔCSH3^ and CrkII^SH2^ constructs showing a similar proportion of dimer, indicating that the SH2 domain mediates dimerization. **c**, Overlaid SEC profiles (S200 10/300) of individually purified CrkII^FL^ dimer and CrkII^FL^ monomer. **d**, Incubation of PEAK1^IDR1^ with CrkII^FL^ dimer (1:1.3 molar ratio) results in formation of a stable complex, as analysed by SEC confirmed by SDS-PAGE analysis of eluted fractions. **e**, SEC-MALS analysis of PEAK2^IDR1^ dimer:CrkII^FL^ dimer complex and PEAK2^IDR1^ dimer alone. Absorbance was measured at a wavelength of 280 nm. Theoretical and observed molecular mass values are indicated. A diagram of the PEAK2^IDR1^ dimer:CrkII^FL^ dimer complex is shown.

**Extended Data Fig. 4:**
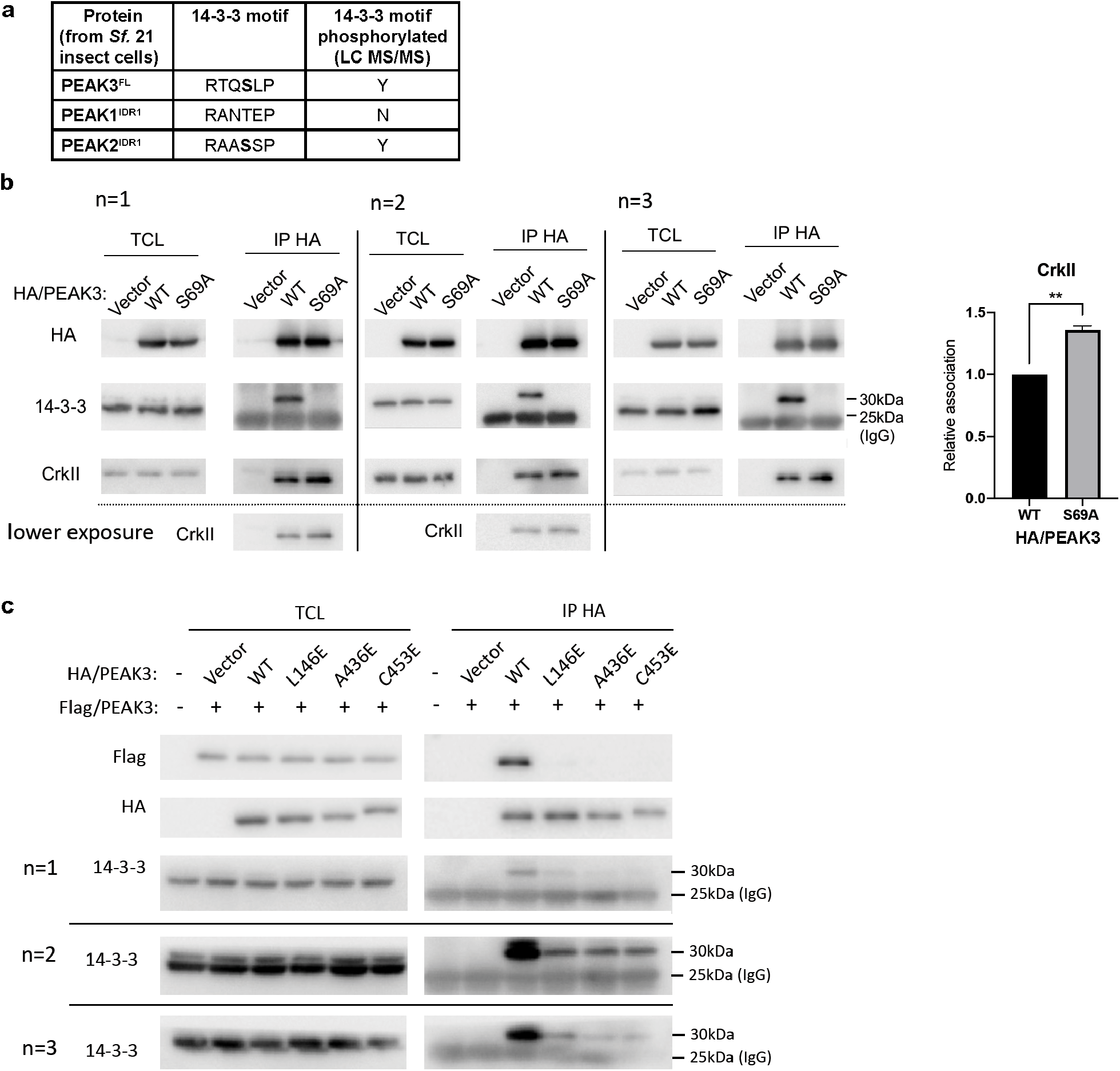
**a**, Table summarising phosphorylation state at the tandem site 14-3-3 motif as identified by LC MS/MS for recombinant PEAK proteins purified from insect cells. Mass-directed tryptic proteomics proteomic analysis identifying phosphorylation sites for recombinantly purified PEAK proteins, as well as identification of 14-3-3 isoforms in the purified PEAK3^FL^:14-3-3 complex purified from insect cells are both available in source data. **b**, Western blot analysis for immunoprecipitation of HA-tagged PEAK3^FL^ WT and PEAK3^FL^ S69A expressed in HEK293 cells and probed using anti-HA, anti-14-3-3 and anti-CrkII; showing that the S69 site is required for 14-3-3 co-immunoprecipitation with PEAK3^FL^ from cells and that mutation of this critical 14-3-3 site (PEAK3^FL^ S69A mutant) is concomitant with a modest increase in the observed level of CrkII bound to PEAK3^FL^. Densitometry data quantifying the mean level relative association of CrkII to HA-PEAK3^FL^ S69A relative to WT. Western blot and densitometry data represent three independent repeats; uncertainties are S.E.M; ** p<0.01, by ratio paired t-test. **c**, PEAK3 mutants (L146E, A436E, C453E) that disrupt dimerization reduce 14-3-3 interaction with PEAK3. Flag-tagged PEAK3 WT and HA-tagged PEAK3 WT or dimerization mutants were co-expressed in HEK293 cells. Anti-HA IPs were prepared from cell lysates and Western blotted as indicated. Data are representative of N=3 independent experiments.

**Extended Data Fig. 5:**
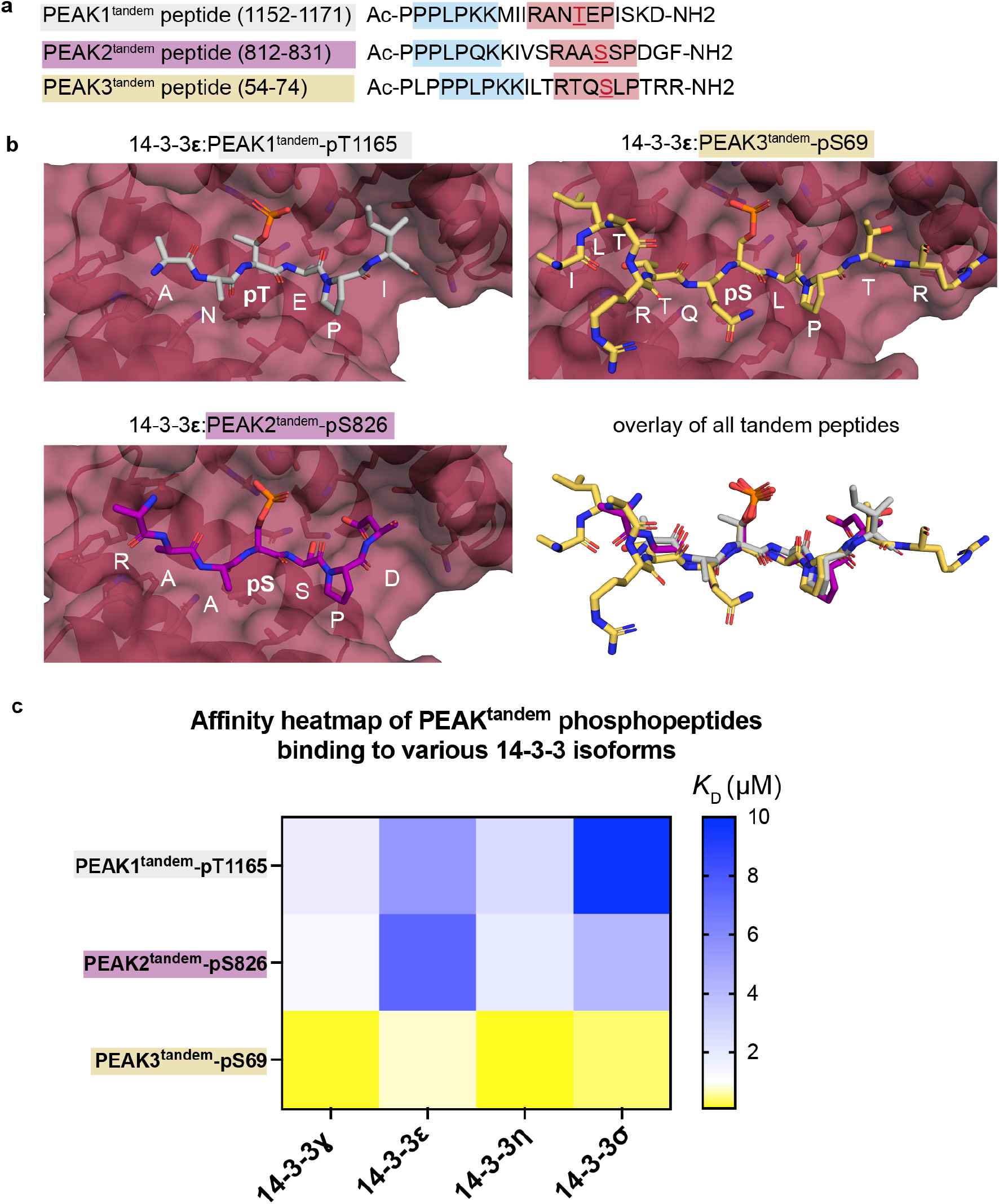
**a**, Primary sequence of PEAK tandem peptides; phosphorylated or non-phosphorylated at the underlined residue (red). **b**, Comparison of crystal structures of PEAK tandem phosphopeptides with 14-3-3ε, showing PEAK1^tandem^-pT1165 (top left, grey sticks), PEAK3^tandem^-pS69 (top right, yellow sticks) and PEAK2^tandem^-pS826 (bottom left, purple sticks) peptides bound to the groove of 14-3-3ε (maroon surface/cartoon) and overlay of all three PEAK peptides (bottom right) demonstrating phospho-recognition by 14-3-3. **c**, Heatmap of SPR steady state affinity values (*K*_D_) of human 14-3-3 isoforms for phosphorylated PEAK tandem peptides (see Supplementary Information for tabulated SPR data).

**Extended Data Fig. 6:**
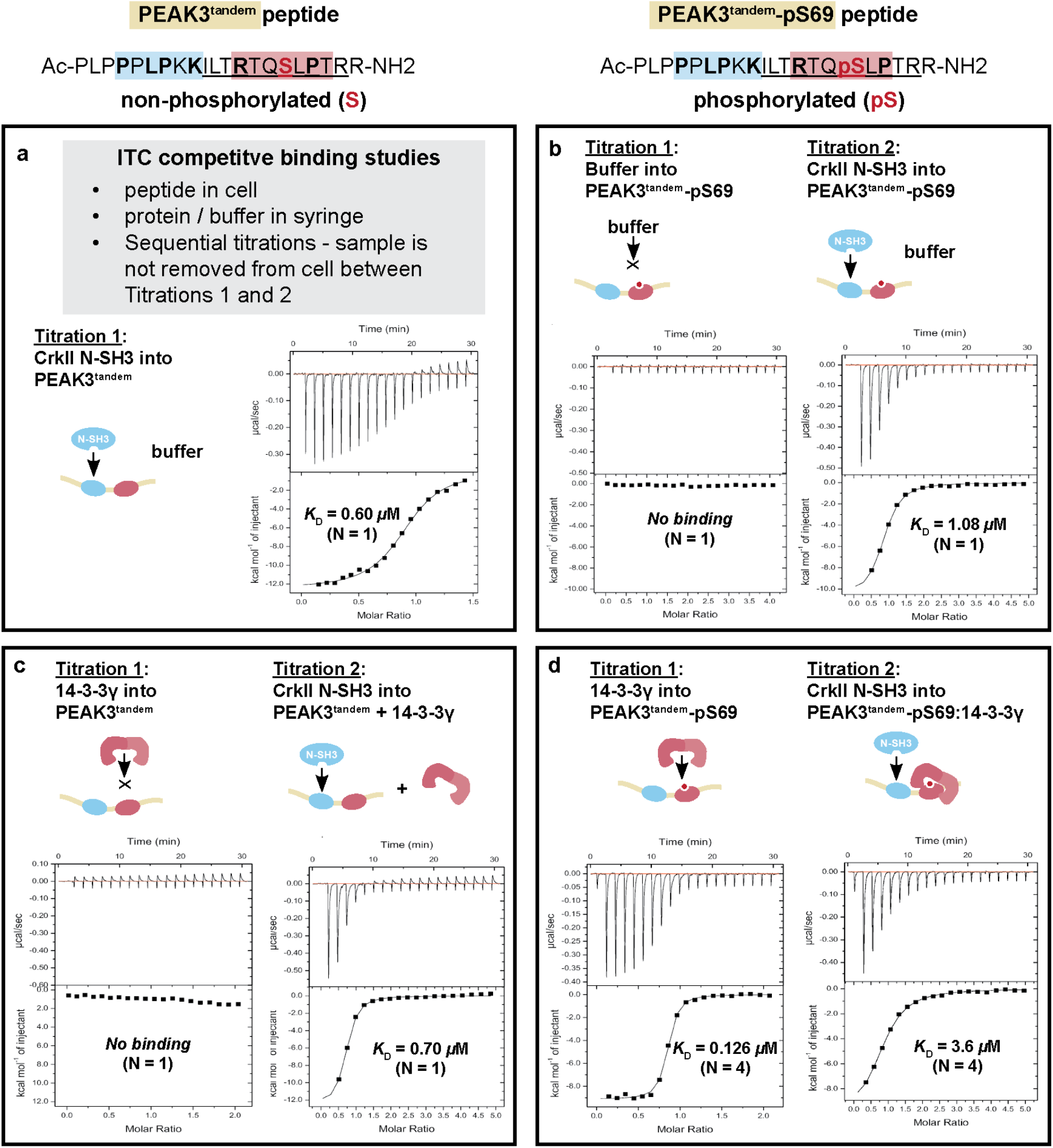
**a-d**, Sequential ITC binding studies using PEAK3^tandem^-S69 (non-phosphorylated, left, a/c) and PEAK3^tandem^-pS69 (phosphorylated, right, b/d) peptides and purified 14-3-3ɣ and CrkII^NSH3^ domain. Corresponding ITC derived affinity values (*K*_D_) (mean value, N > 1) and the number of independent experiments (N) are shown on the representative plot for each titration (see Supplementary Information for tabulated ITC data).

**Extended Data Fig. 7:**
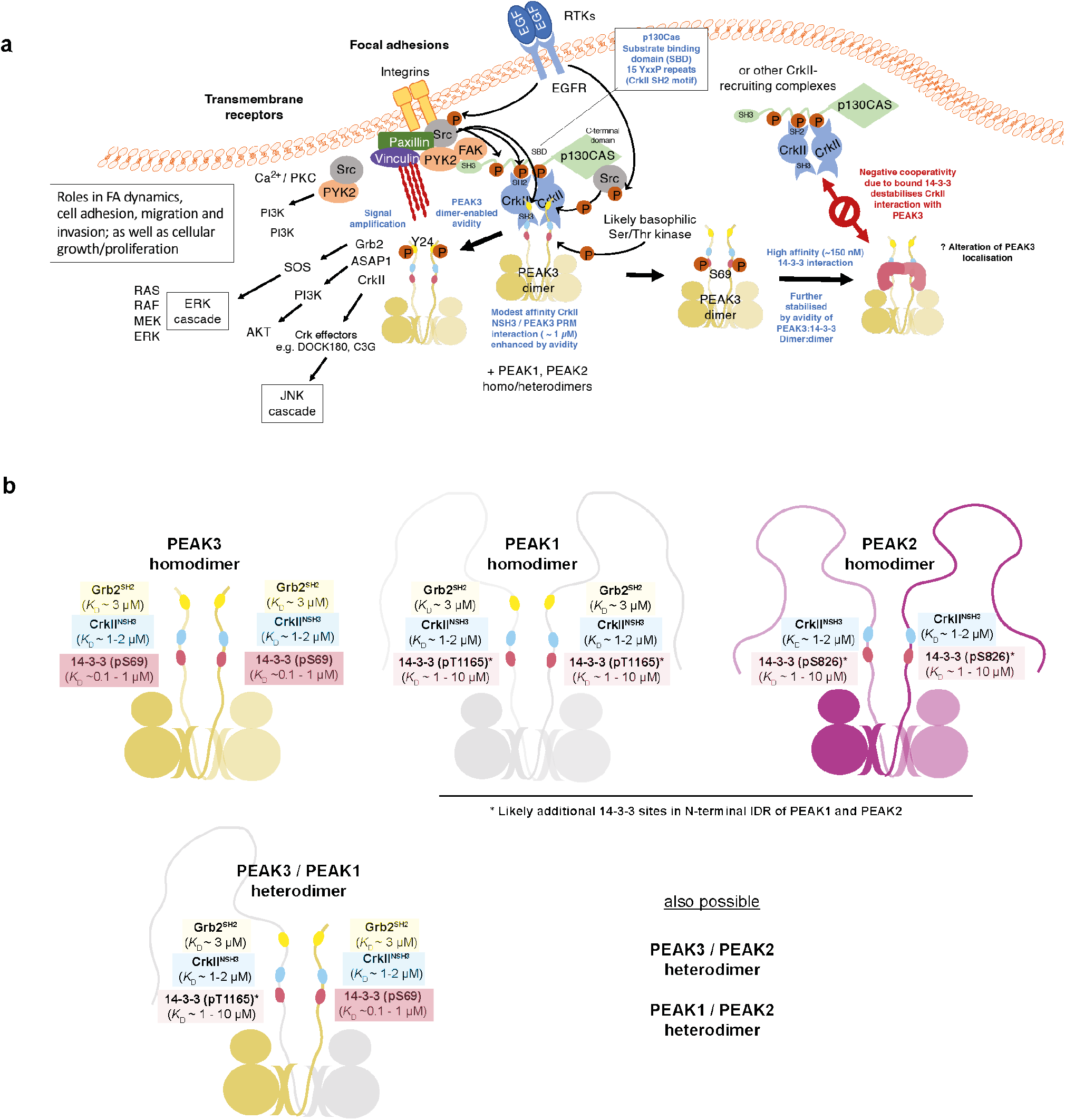
**a**, Schematic of PEAK signaling pathways at focal adhesions. **b**, Schematic summarizing the approximate binding affinity (*K*_D_ for 1:1 interaction) of Grb2, CrkII and 14-3-3 towards PEAK1/PEAK2/PEAK3 dimers at N-terminal IDR motifs examined in this study, highlighting PEAK homodimer and heterodimer configurations and diverse interaction outputs.

