## Supplementary material for "When two’s a crowd - Structural mapping of PEAK pseudokinase interactions identifies 14-3-3 as a molecular switch for PEAK3/Crk signaling": Source Data

**Source data to Fig. 2: MS/MS analysis of tryptic peptides following *in vitro* kinase assay of PEAK3-FL and PEAK1^IDR1^ purified from insect cells.** **a,** (i) PEAK3 control sample is not phosphorylated at Y24 (SH2 motif) in the absence of Src or Abl. (ii) PEAK3 is phosphorylated at Y24 (SH2 motif) upon incubation with Src. (iii) PEAK3 is not phosphorylated at Y24 (SH2 motif) upon incubation with Abl.  **b,** (i) PEAK1^IDR1^ control sample is not phosphorylated at T1007 (SH2 motif) in the absence of Src or Abl. (ii) PEAK1^IDR1^ is phosphorylated at T1007 (SH2 motif) upon incubation with Src. (iii) PEAK1^IDR1^ is not phosphorylated at T1007 (SH2 motif) upon incubation with Abl.  **c,** Positive controls demonstrating Abl and Src *in vitro* kinase activity towards CrkII-FL. (i) CrkII^FL^ control sample is not phosphorylated at Y221 in the absence of Src or Abl. (ii) CrkII^FL^ is phosphorylated at Y221 (known SFK site) and Y upon incubation with Src. (iii) CrkII^FL^ is phosphorylated at Y221 (known SFK site) upon incubation with Abl.

a. ***in vitro* kinase assay of recombinant PEAK3-FL**

**(i) PEAK3 control sample is not phosphorylated at Y24 (SH2 motif) in the absence of Src or Abl.**

PTWSTQTYSNLGQIR

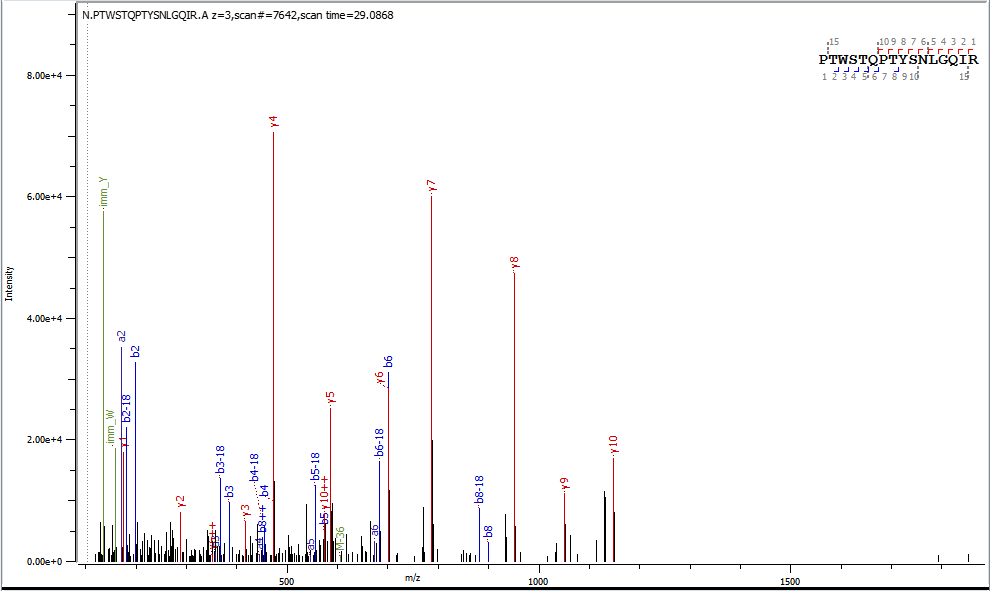

**(ii) PEAK3 is phosphorylated at Y24 (SH2 motif) upon incubation with Src**

PTWSTQPT**Y_p24_**SNLGQIR

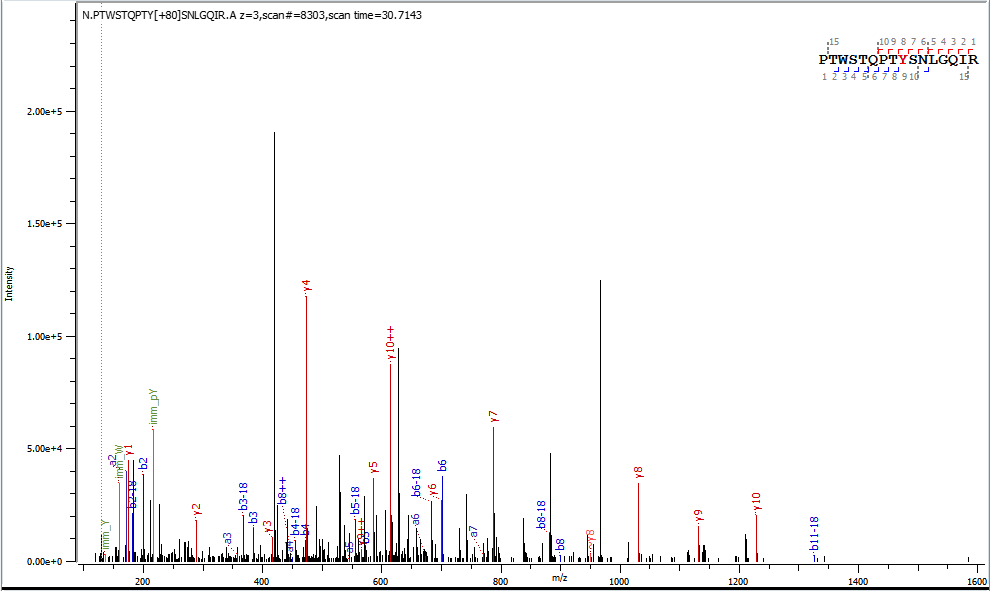

**(iii) PEAK3 is not phosphorylated at Y24 (SH2 motif) upon incubation with Abl.**

PTWSTQPTYSNLGQIR

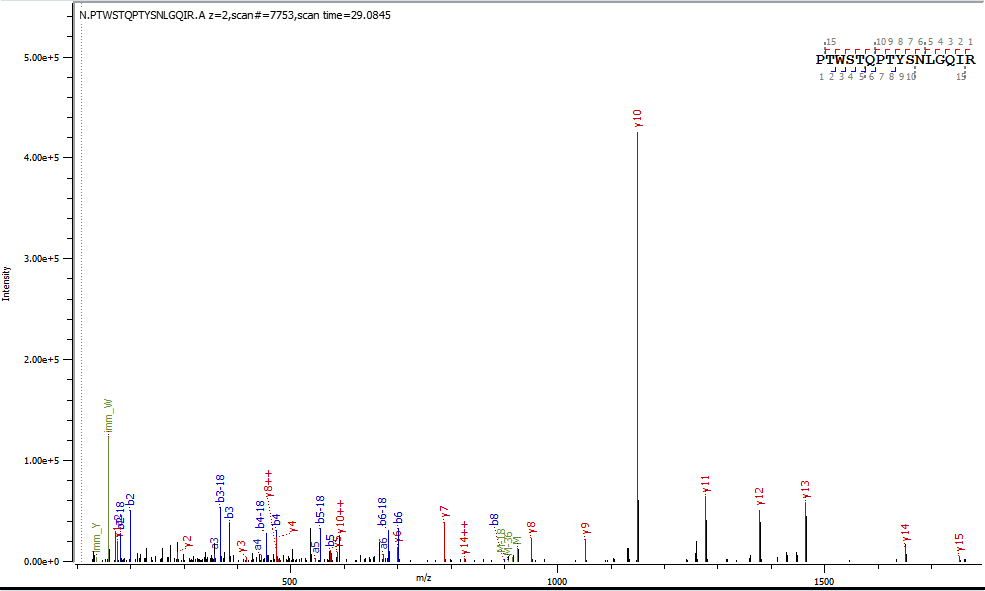

b. ***In vitro* kinase assay - PEAK1^IDR1^ (*Sf21*)**
(i) **PEAK1^IDR1^ control sample is not phosphorylated at T1007 (SH2 motif) in the absence of Src or Abl**

EDGKEDISDPMDPNPCSATYSNLGQSR

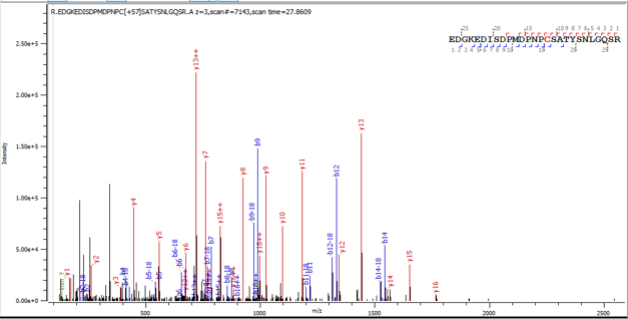

(ii) **PEAK1^IDR1^ is phosphorylated at T1007 (SH2 motif) upon incubation with Src**

EDGKEDISDPMDPNPCSATY_p1007_SNLGQSR

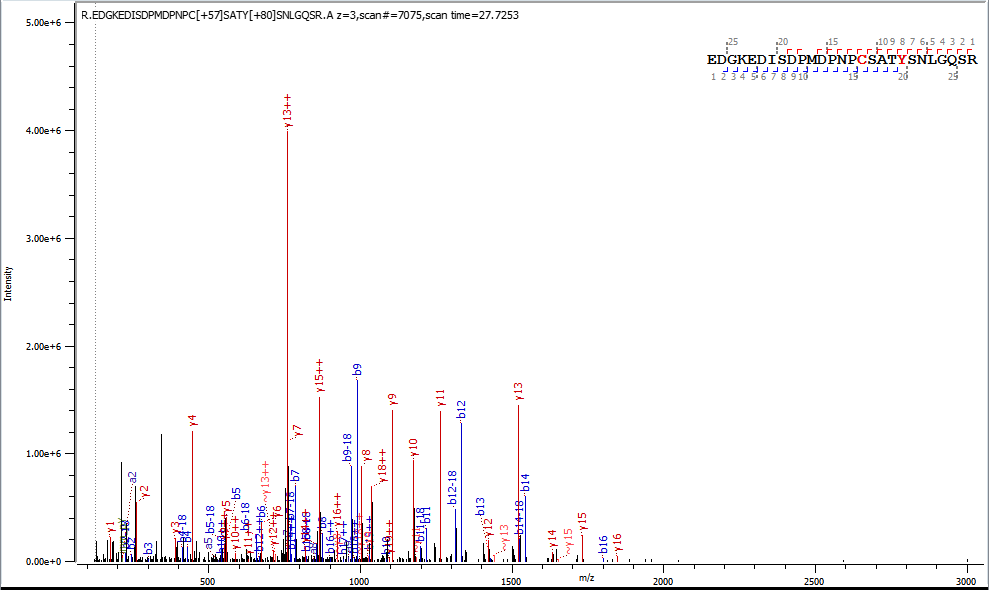

(iii) **PEAK1^IDR1^ is not phosphorylated at T1007 (SH2 motif) upon incubation with Abl.**

SATYSNLGQSR

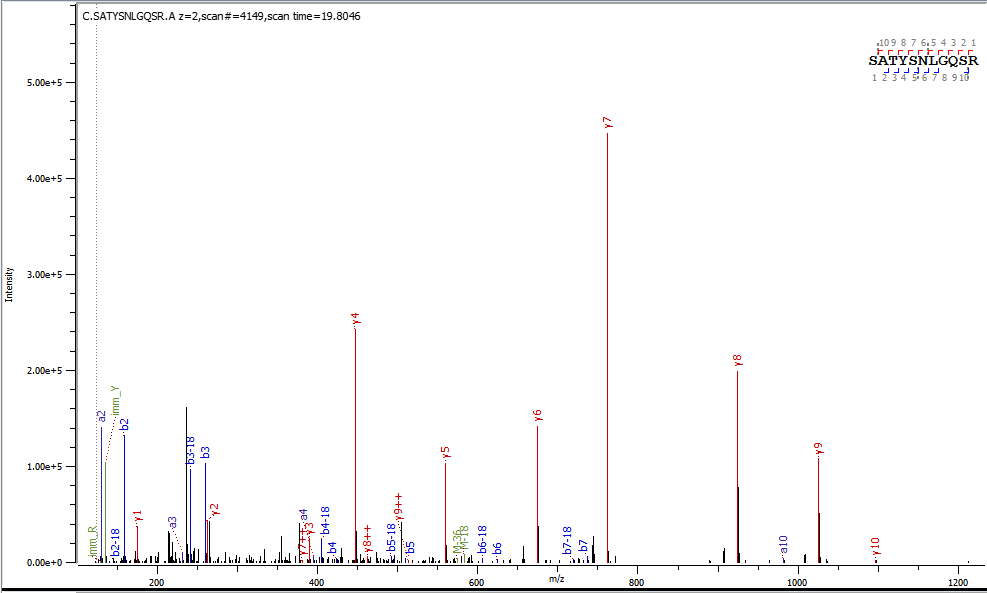

c. ***In vitro* kinase assay – CrkII^FL^ monomer (*E.coli*) – positive control**
(i) **CrkII^FL^ control sample is not phosphorylated at Y221 in the absence of Src or Abl**

YRPASASVSALIGGNQEGSHPQPLGGPEPGPYAQPSVNTPLPNLQNGPIYAR

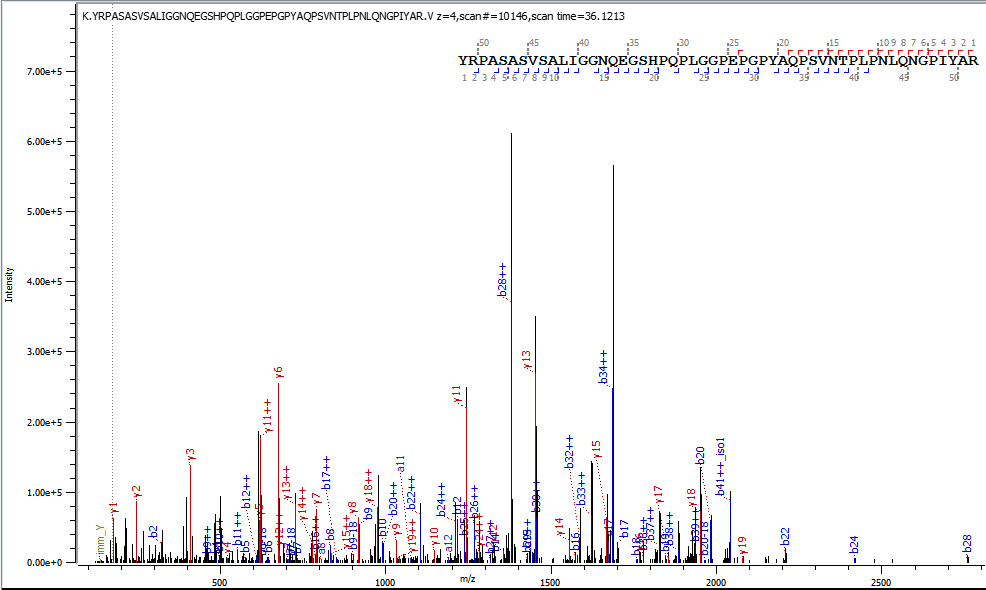

(ii) **CrkII^FL^ is phosphorylated at Y221 (known SFK site) upon incubation with Src**

YRPASASVSALIGGNQEGSHPQPLGGPEPGPY_p221_AQPSVNTPLPN

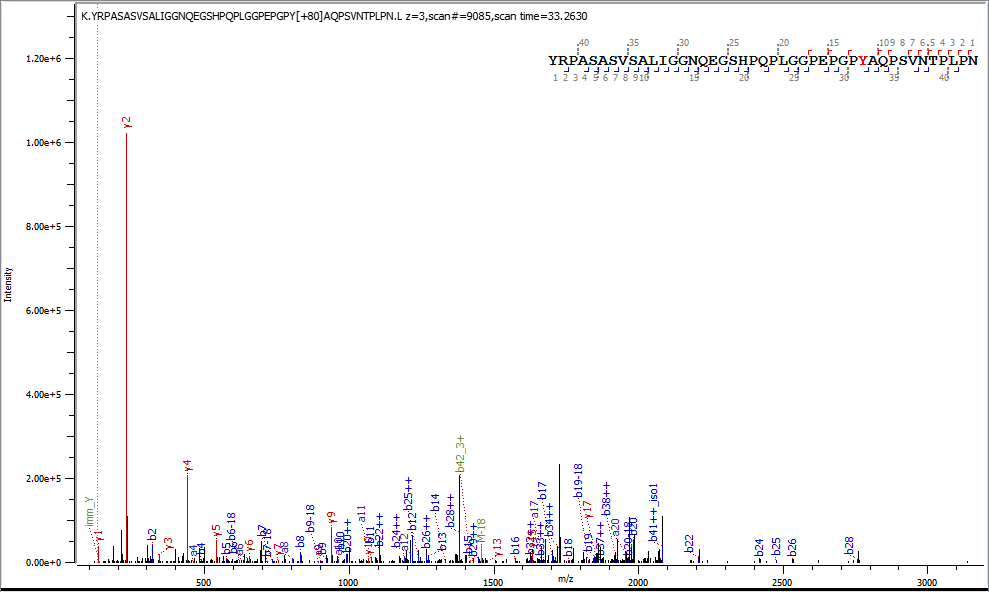

(iii) **CrkII^FL^ is phosphorylated at Y221 (known SFK site) upon incubation with Abl**

YRPASASVSALIGGNQEGSHPQPLGGPEPGPY_p221_AQPSVNTPLPNL

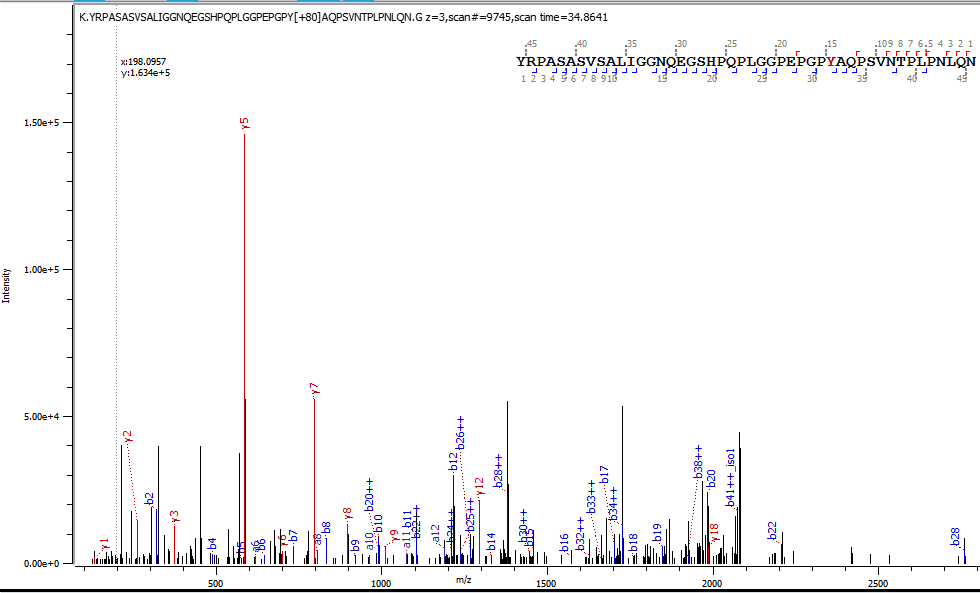

**Source Data to Fig. 4a. (i),** Mass-directed tryptic proteomics showing identification of 14-3-3ε and 14-3-3ζ isoforms as PEAK3 interactors. **(ii),** LC-MS/MS analysis of recombinant PEAK3 showing that PEAK3 14-3-3 site is phosphorylated at S69. **(iii),** LC-MS/MS analysis of recombinant insect cell expressed PEAK2^IDR1^ showing that it is phosphorylated at S826/S827 within the 14-3-3 motif (tandem site). **(iv),** LC-MS/MS analysis of recombinant insect cell expressed PEAK1^IDR1^ showing that it is not phosphorylated at the 14-3-3 motif (tandem site).

(i) **Mass-directed tryptic proteomics showing identification of 14-3-3ε,ζ isoforms**

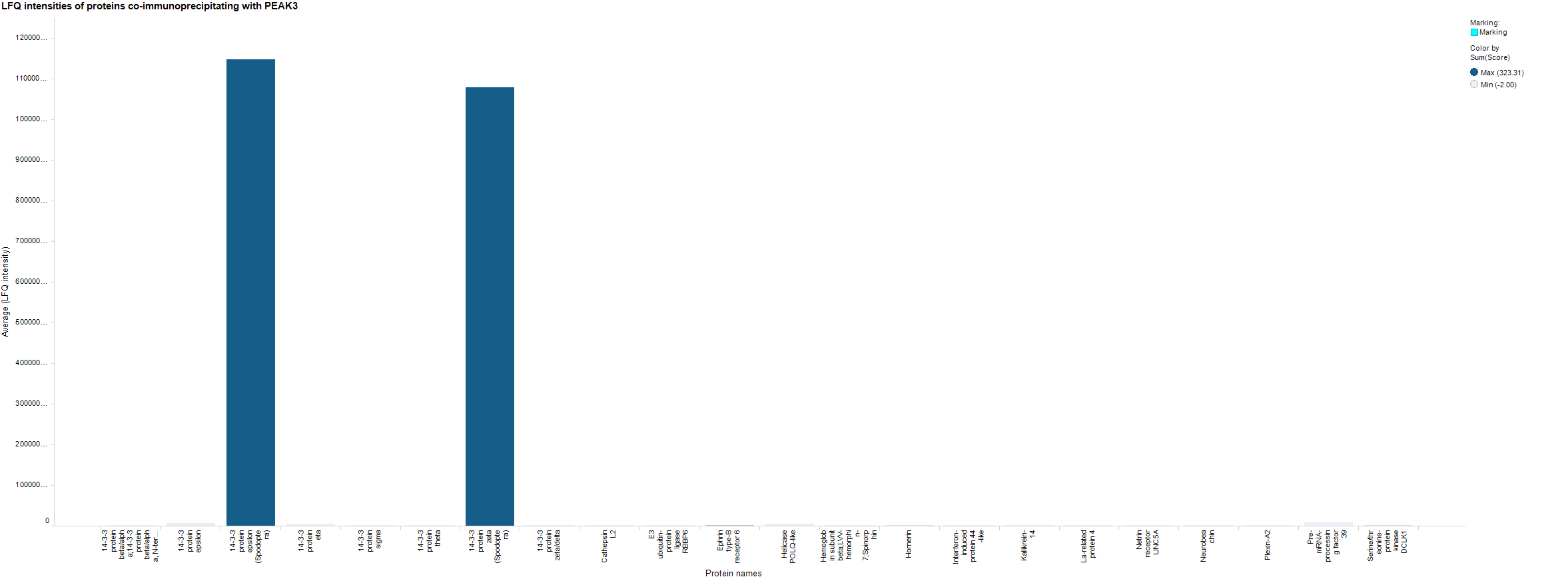

(ii) **PEAK3 14-3-3 site is phosphorylated at S69 within the 14-3-3 motif (tandem site)**

**TQS_p69_LPTRR**

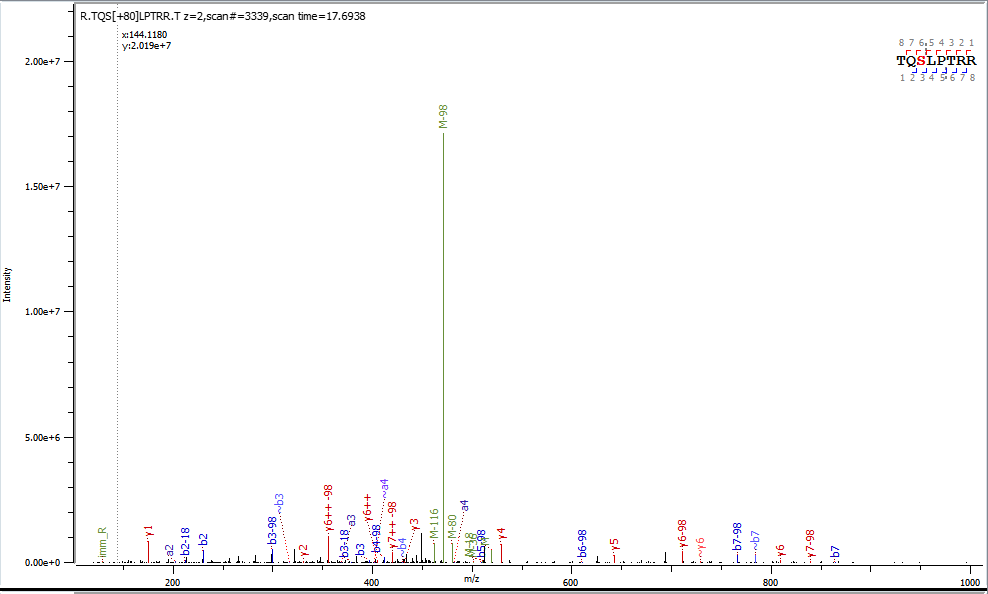

(iii) **PEAK2^IDR1^ is phosphorylated at S826/S827 within the 14-3-3 motif (tandem site)**

**AAS_p826_SPDGFFWT**

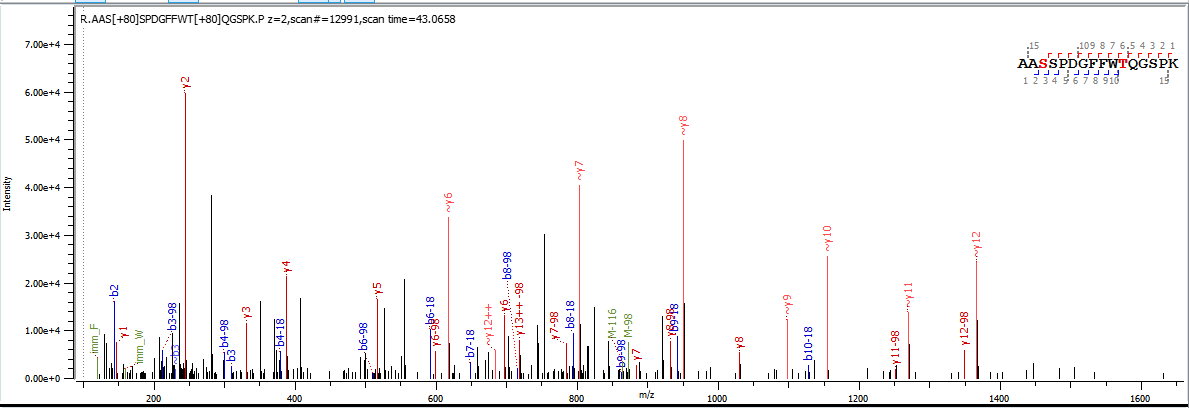

**AASS_p827_PDGFFWT**

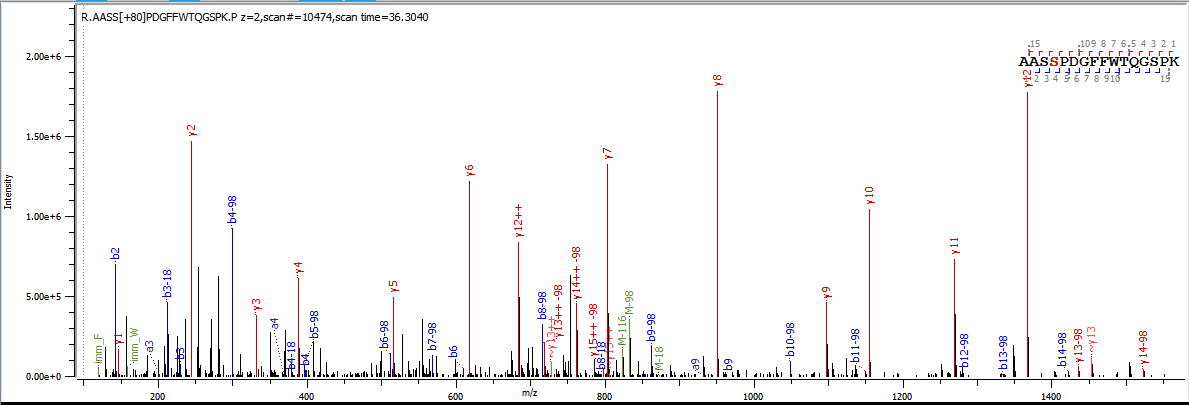

(iv) **PEAK1^IDR1^ is not phosphorylated within the 14-3-3 motif (tandem site)**

**ANTEPISKDLQK**

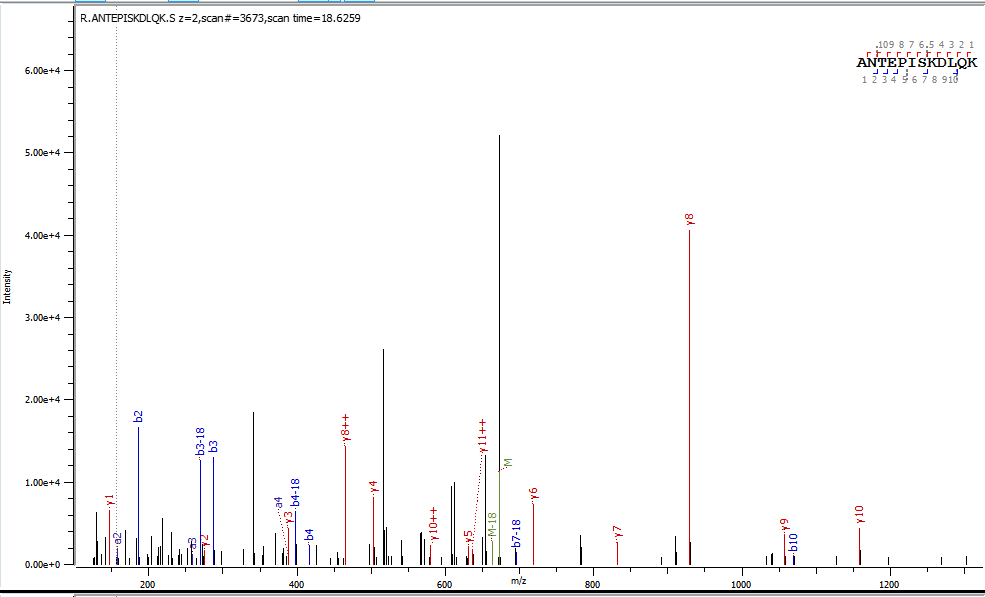

**Source Data to Fig. 4b and Extended Fig. 4b. Uncropped gels**

**14-3-3**

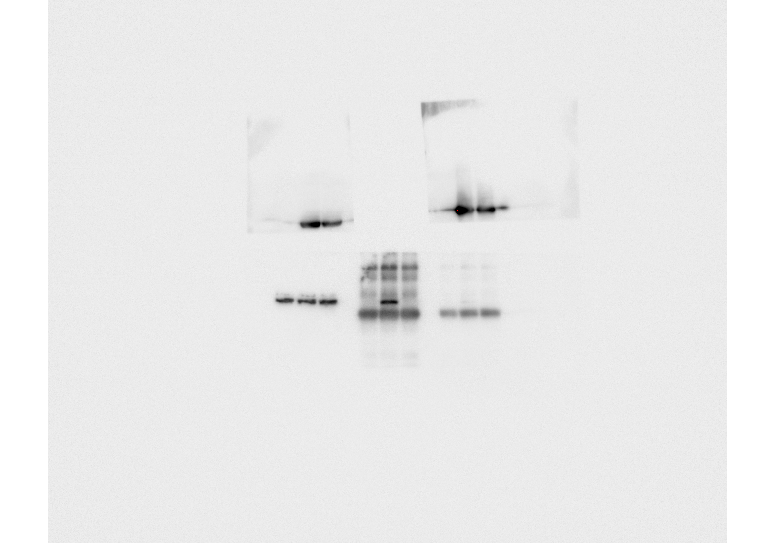

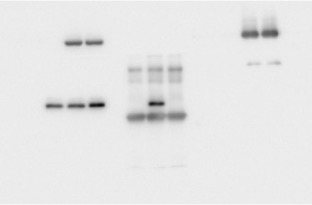

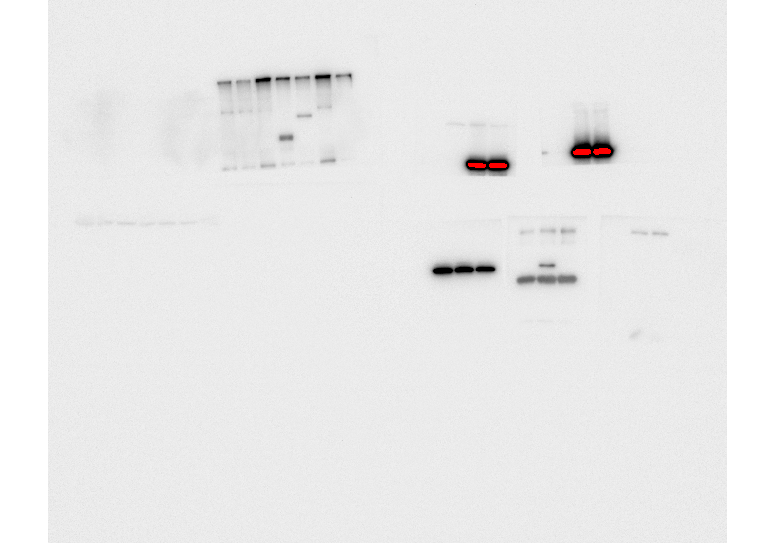

n=2

n=3

n=2

n=1

**Source Data to Extended Data Fig. 4b. Uncropped gels**

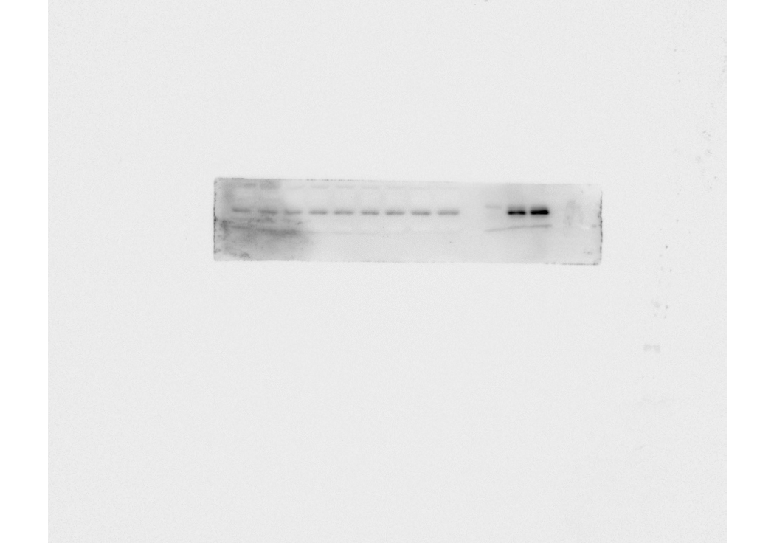
**CrkII**

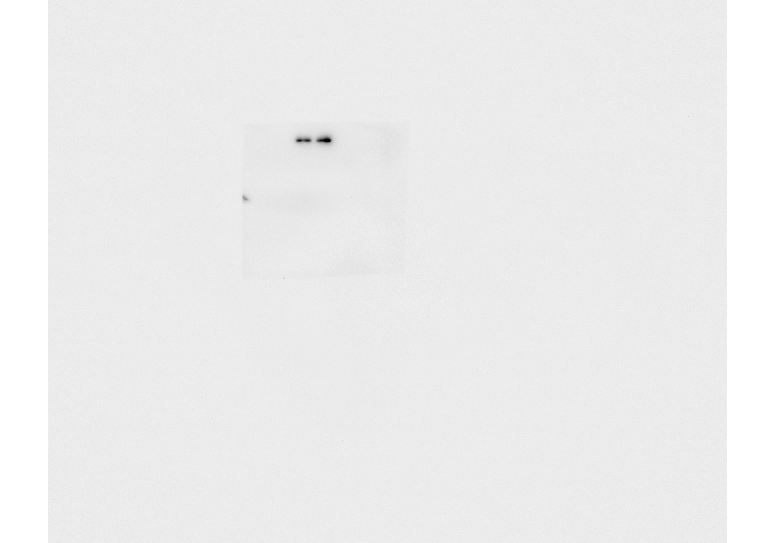

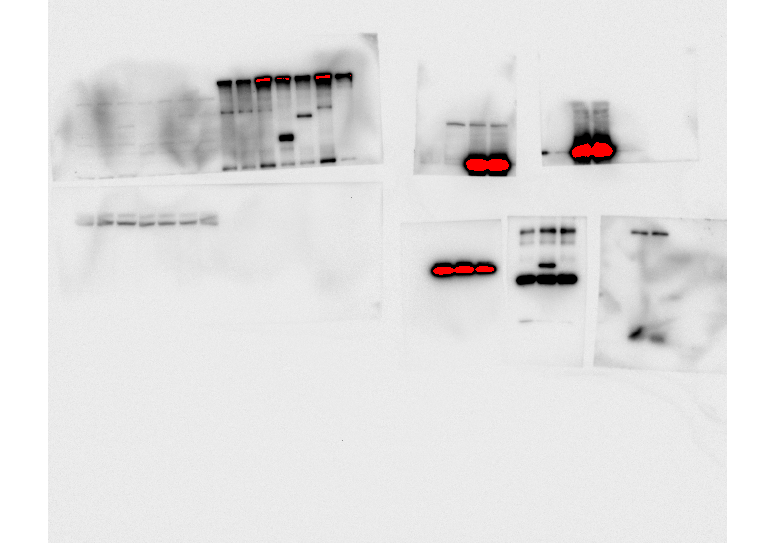

n=1

n=3

n=2

**Source Data to Extended Data Fig. 4c. Uncropped gels**

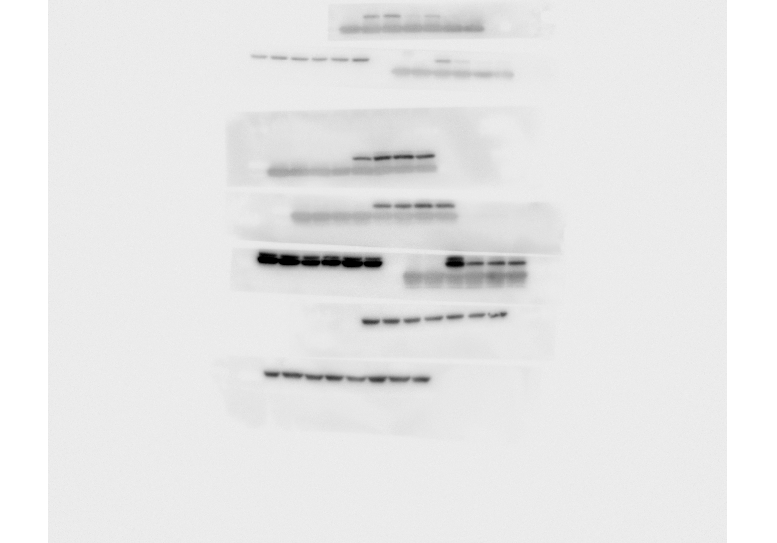

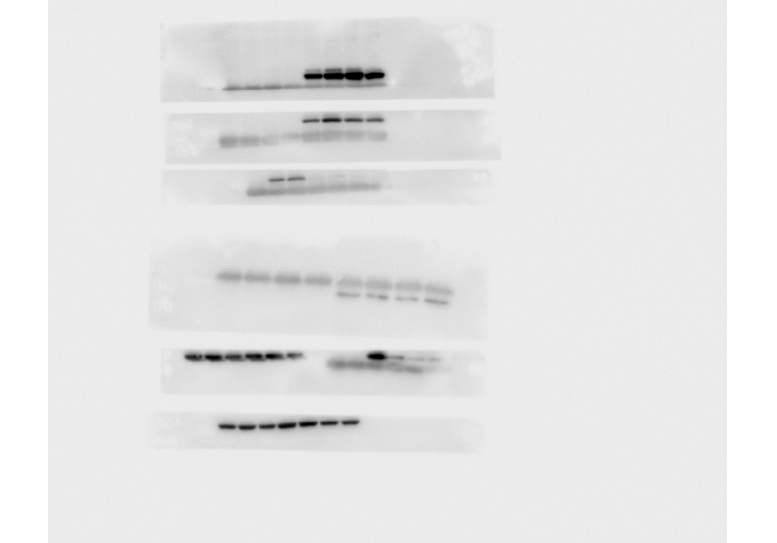

n=3
